# Nowhere to Hide: The Effect of Centromere Architecture on LTR Retrotransposon Dynamics

**DOI:** 10.64898/2026.07.14.738416

**Authors:** Marie Krátká, Klára Panda, Pavel Jedlička, Petr Bureš, Zdeněk Kubát, Jakub Šmerda, André Marques, Eduard Kejnovský, František Zedek

**Author notes:** shared first authors (Marie Krátká, Klára Panda).

## Abstract

Centromere architecture can determine where transposable elements persist, yet its effects on retrotransposon turnover remain poorly understood. Here we test a chromosome-level “nowhere-to-hide” model, in which holocentric chromosomes, owing to distributed centromere activity and reduced chromatin compartmentalization, provide fewer stable repeat-rich refugia than monocentric chromosomes. We combined genome-wide LTR retrotransposon annotation, spatial modelling and FISH across 40 holocentric plant species and 31 closely related monocentric relatives from Poales, *Cuscuta*, and Melanthiaceae. Overall LTR retrotransposon abundance and Ty1-copia/Ty3-gypsy composition were explained mainly by lineage history and chromosome size, rather than by holocentricity itself. By contrast, element persistence and removal showed dependence on centromere architecture. Intact LTR retrotransposons were younger in holocentric genomes, and holocentric chromosomes lacked the chromosome-size-dependent spatial clustering of element age observed in monocentrics. Solo-LTR profiles further revealed weaker spatial clustering of removal signatures in holocentric chromosomes, consistent with a more homogeneous chromosome-wide landscape of ectopic recombination. Epigenomic analyses of a matched *Luzula*–*Juncus* pair indicated that young elements can occur in centromeric chromatin, whereas solo LTRs are associated with more euchromatic contexts. These results support the nowhere-to-hide model, showing that centromere architecture does not shape LTR retrotransposons accumulation, but their persistence and removal efficiency.

## Introduction

Transposable elements (TEs) constitute a major proportion of plant genomes and are key drivers of genome evolution, structure, and gene regulation (Bennetzen et al., 2005; Biémont & Vieira, 2006; Kejnovsky et al., 2012; Wells and Feschotte, 2020). Among TEs, long terminal repeat retrotransposons (LTR-RTs) are the most abundant in plant genomes (Kumar & Bennetzen, 1999) and may constitute up to 85% of the nuclear DNA in large-genome species, such as cereals (Schnable et al., 2009; Wicker et al., 2018).

LTR-RTs propagate through a “copy-and-paste” mechanism, in which reverse transcription of an RNA intermediate and subsequent integration into the genome lead to their amplification (Kumar and Bennetzen, 1999; Wicker et al., 2007). LTR-RT proliferation often occurs in episodic bursts of retrotransposition, triggered by stress, hybridization, or polyploidization (Charles et al., 2008; Makarevitch et al., 2015; Horváth et al., 2017). Because the two LTRs are identical at the time of insertion, divergence between LTR ends of the same element can be used to estimate insertion age (Jedlicka et al., 2020). LTR-RT accumulation is counterbalanced by two ectopic recombination-based removal processes (Devos et al., 2002; Morales-Díaz et al., 2025). First, homologous unequal recombination between the two LTRs of the same or two distinct elements can excise the whole region between the recombining LTRs, leaving only one long terminal repeat behind (solo LTR) as a removal signature. Second, illegitimate recombination through double-strand break misrepair produces deletions and truncated copies of LTR-RTs. Together, these ectopic recombinations contribute to reducing the genomic load of retrotransposons and generate abundant solo LTRs and/or fragmented elements (Devos et al., 2002; Kent et al., 2017; Jedlicka et al., 2020; Morales-Díaz et al., 2025).

TE turnover – balance between LTR-RT amplification and removal – plays a major role in shaping genome size (Devos et al., 2002; Bennetzen et al., 2005; Kejnovsky and Jedlicka, 2022), and may even facilitate speciation by altering genome architecture and gene regulation (Oliver & Greene, 2009; Rebollo et al., 2010). Most insertions are considered deleterious (Deniz et al., 2019), and their activity is therefore suppressed by epigenetic silencing mechanisms, such as DNA methylation, histone modifications, and RNA interference, including the action of small RNA molecules and post-transcriptional gene silencing (Kejnovsky and Jedlicka, 2022), or by ectopic recombination removal. TEs preferentially accumulate in low-recombining areas such as (peri)centromeres and other heterochromatic clusters (Schnable et al., 2009), as well as in non-recombining regions like the Y chromosomes (Kejnovsky et al., 2009; Wang et al., 2014; Hobza et al., 2017), where their removal is less efficient. In addition to intra-chromosomal position, the genome-wide recombination rate (cM/Mb) scales negatively with chromosome length, meaning that larger chromosomes generally experience lower recombination densities (Lynch et al., 2011; Brazier & Glémin, 2022; Zedek et al., 2026; Zhang et al., 2026a). Although the association between meiotic and ectopic recombination has been established mainly in non-plant systems (Lichten et al., 1987; Myers et al., 2008; McVean, 2010; Kent et al., 2017), unequal recombination also contributes to LTR-RT removal in plants (Ma et al., 2004; Cossu et al., 2017). Extrapolating from these observations, larger chromosomes may have a reduced capacity for LTR-RT removal and may consequently act as large-scale genomic sinks for repetitive DNA accumulation, irrespective of centromere position.

This pattern has been observed in species with monocentric chromosomes, which possess a single centromeric region. There, chromatin is typically compartmentalized, with euchromatic domains characterized by higher recombination rates and heterochromatic clusters displaying recombination suppression (Kent et al., 2017; Wang et al., 2024). Such architecture provides genomic niches where LTR retrotransposons and other repetitive elements can accumulate and persist over extended evolutionary timescales. In contrast, holocentric chromosomes, which assemble kinetochores along their entire length (Bureš et al., 2013; Mandrioli and Manicardi, 2020; Hofstatter et al., 2021; Marques and Drinnenberg et al., 2025), lack a localized centromere and are therefore expected to differ fundamentally in their recombination landscape. It has been proposed that holocentric chromosomes do not provide stable, recombination-suppressed refugia for repetitive DNA, effectively leaving transposable elements with “nowhere to hide” from selection and recombination-mediated removal (Bureš & Zedek, 2014). Consistent with this idea, holocentric genomes often exhibit decompartmentalized chromatin lacking clear euchromatic and heterochromatic clusters (Mandrioli & Manicardi, 2012; Heckmann et al., 2013; Hofstatter et al., 2022; Mata-Sucre et al., 2024), and meiotic recombination may be largely decoupled from local sequence or epigenetic state (Zhang et al., 2026b, Haenel et al., 2018; Shipilina et al., 2022; Castellani et al., 2024). However, the effect of decompartmentalized chromatin in holocentrics on distribution and overall rate of TE removal has not been directly tested. Additionally, the mechanistic basis for the previously reported correlation between meiotic crossover frequency and unequal homologous recombination rates remains unresolved. This might affect the TE removal rate, since holocentricity is associated with a global decline of cross-over frequency (Zedek et al., 2026).

Here we aim to test these four predictions: (i) if holocentric chromosomes indeed lack recombination-suppressed refugia, the distribution of LTR-RT removal signatures should be more even along their length compared to monocentric chromosomes, measurable as a more uniform distribution of ratio of solo LTRs to full-length LTR-RTs; (ii) LTR-RT insertions should be, on average, younger in holocentric than in monocentric species, where elements can persist longer in heterochromatic regions; (iii) the distribution of insertion ages is also expected to be more homogeneous along holocentric than monocentric chromosomes; and (iv) increasing chromosome size will generally reduce removal efficiency in both systems, but holocentric chromosomes will maintain a younger LTR-RT age structure than monocentrics across the entire size spectrum due to their distributed recombination landscape

To address these predictions, we combined genome-wide bioinformatic annotation with cytogenetic visualization (FISH). Holocentricity has been reported in several angiosperm lineages: family Cyperaceae (Poales), and genera *Chionographis* (Melanthiaceae, Liliales), *Cuscuta* (Convolvulaceae, Solanales), *Drosera* (Droseraceae, Caryophyllales), *Luzula* (Juncaceae, Poales), *Morus* (Moraceae, Rosales), and *Myristica* (Myristicaceae, Magnoliales; Marques and Drinnenberg, 2025). From these lineages, we focused on Poales, *Cuscuta*, and Melanthiaceae (**Fig. 1**), which currently provide closely related monocentric and holocentric taxa with high-quality chromosome-level assemblies, allowing us to analyze how centromere organization and chromosome size shape LTR retrotransposon distribution, age structure, and epigenetic regulation across entire chromosomes.

**Figure 1.**
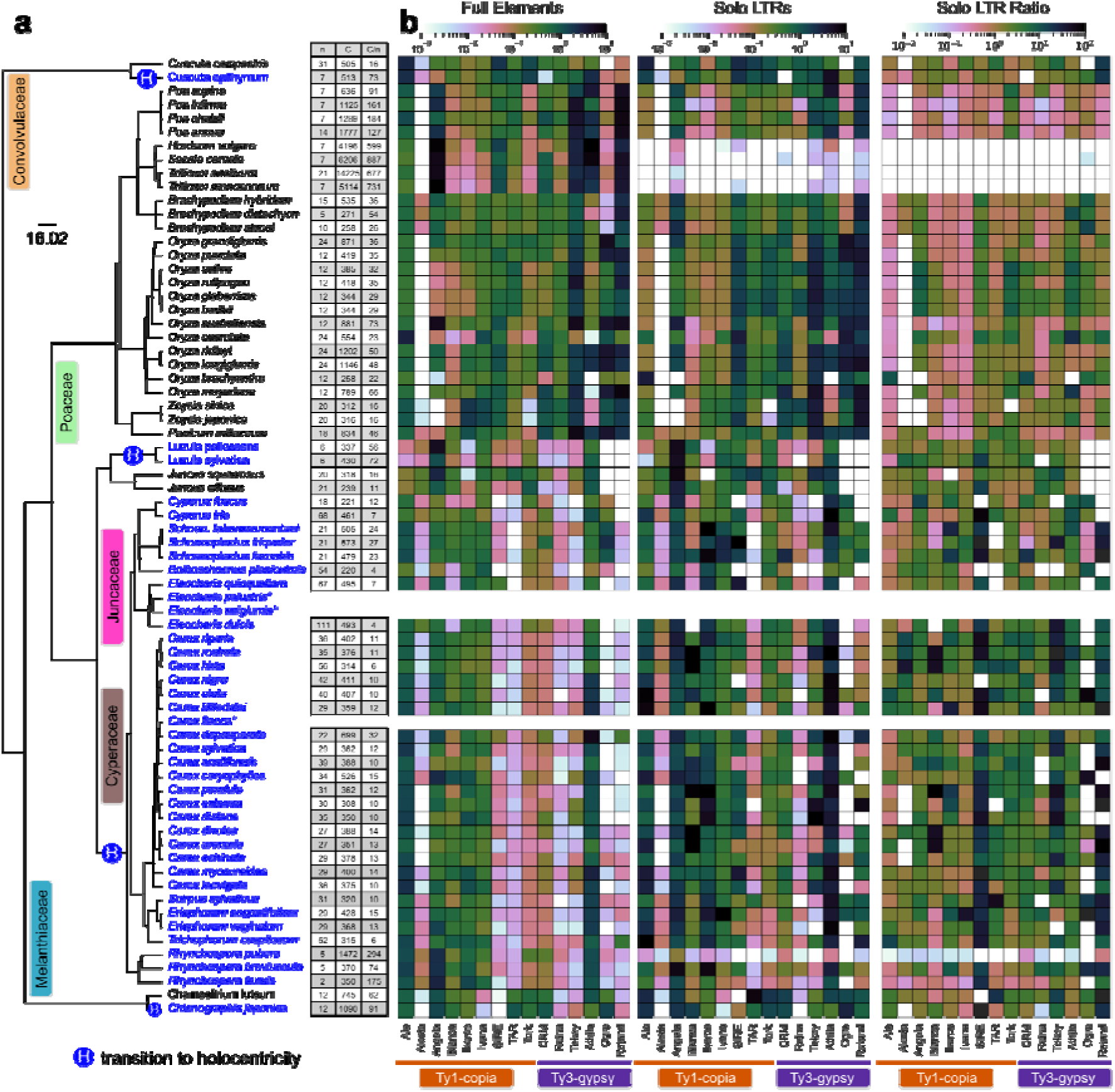
Included species and LTR-RT abundance. **(a)** Four groups where the transition to holocentricity (branches marked by H) has occurred are included in the analysis—Cyperaceae, Juncaceae, Convolvulaceae, and Melanthiaceae. For each species in the bioinformatics analysis, haploid chromosome number (n), total chromosome-level assembly size in Mbp (C), and average chromosome size (C/n) are reported (**Supplementary Table S2**). Asterisk (*) indicates species used only for the cytogenetic analysis. Two subspecies of *El. uniglumis* (ssp. *uniglumis* and ssp. *sterneri*) were used in the cytogenetic analysis (**Supplementary Table S3**). **(b)** Heatmap panels show the abundance of annotated full-length elements and solo LTRs as the number of elements per 1 Mbp and the ratio of solo LTR to full-length elements. The plotted values are log10 transformed with color bar ticks back-transformed to match the original values. The element abundance profile is phylogenetically clustered.

## Material and methods

All analyses were performed using open-source tools (versions and references are listed in **Supplementary Table S1**). Custom scripts and pipelines are available at [https://github.com/437364/nowhere_to_hide_LTR-RTs].

### Species selection

We have used genome assemblies available from the National Center for Biotechnology Information (NCBI; **Supplementary Table S2**). Our target group were holocentric plants with chromosome-level or complete genome assemblies available at the time of analysis (August 2025). To provide a comparative framework, we also included closely related monocentric species from the same or closely related families. The monocentric dataset was selected to cover a broad range of genome sizes and chromosome numbers and to achieve a similar number of monocentric and holocentric species. To represent phylogenetic relationships between the selected species, we pruned a species-level angiosperm tree (Forest, 2023) using a custom script and visualized the resulting tree using the phylo.io (Robinson et al., 2016; **Supplementary Table S1**).

Assembly quality was assessed with BUSCO (Seppey, 2019) against the Viridiplantae odb10 dataset to assess gene completeness, and gap density was estimated using seqtk gap (**Supplementary Table S1**). Results were visualized using a custom script (see https://github.com/437364/nowhere_to_hide_LTR-RTs). Since the availability of high-quality assemblies is currently limited for holocentric Cyperid species with large chromosomes, we complemented the bioinformatic dataset with FISH-based analyses of selected species.

### Genome annotation

We developed a pipeline for comprehensive annotation of LTR-RTs. First, LTR-RTs were annotated using the DANTE-LTR pipeline with parameters set to tolerate three missing domains (-M 3; Novák et al. 2024). This pipeline annotates LTR-RTs based on both protein domains and structural features. Identified elements were classified according to their DANTE_LTR rank as full-length elements, when both protein domains and LTRs were detected, or partial elements, when protein domains were detected but complete element structure was not recovered. In four Poaceae species (*H*. *vulgare*, *Se. cereale*, *Trit. aestivum*, and *Trit. monococcum*), large chromosome size caused processing issues. To avoid an excessive demand for computational resources, only one or two chromosomes from each of these species were included in the analysis (**Supplementary Table S2**).

Full-length LTR-RTs were used to build a repeat library with the dante_ltr_to_library command. This library was then used for a similarity-based search using RepeatMasker (Smit et al., 2013–2015, **Supplementary Table S1**). Solo LTRs were identified using a script from Ou and Jiang (2018; **Supplementary Table S1**), based on the absence of nearby internal LTR-RT region from the same family and strand.

Genomic distributions of LTR-RT features were analysed in sliding windows along chromosomes using a window size of 5% of chromosome length and a step size of 1%. For each window, we calculated the abundance of full-length elements, partial elements, solo LTRs, and solo LTR ratio.

Solo LTR ratio was calculated as the number of solo LTRs divided by the number of intact full-length LTR-RTs from the same window, chromosome, species, or LTR-RT family, depending on the analysis. This definition follows common usage in published LTR-RT studies and is reported throughout the manuscript as the solo/intact LTR-RT ratio. For GLMMs, the ratio was log-transformed and window-level models were weighted by the number of LTR-RT elements contributing to each ratio.

### Spatial pattern analyses

Spatial patterns along chromosomes were quantified using Moran’s index (Moran, 1950) and Dynamic Time Warping (DTW) clustering. Moran’s I provides a single spatial autocorrelation metric for the whole chromosome, where values near zero indicate spatial independence and high positive values show spatial clustering. DTW clustering captures the shape of the pattern. Chromosomes were divided into sliding windows of 5% of chromosome length with a 1% step size. Moran’s I was calculated separately for window-level LTR identity and for the proportional solo LTR ratio. The main analyses used a neighbourhood threshold of 0.2, corresponding to 20% of chromosome length, as an intermediate chromosomal scale intended to capture broad regional clustering while avoiding both very local window-to-window noise and near-global comparisons across large chromosome fractions. Additional threshold-sensitivity analyses were performed across a range of neighbourhood thresholds (0.05, 0.1, 0.3, 0.4, 0.5, and 0.6) to assess how spatial autocorrelation changed with scale.

For DTW clustering, we used the same window-based chromosome profiles to characterize spatial distribution archetypes. Distribution profiles of mean LTR identity, and proportional solo LTR ratio were analyzed. Profiles with low LTR-RT content were excluded according to criteria reported in [https://github.com/437364/nowhere_to_hide_LTR-RTs]. Pairwise distances between profiles were calculated using Dynamic Time Warping function implemented in the dtaidistance Python library to account for variable monocentromere position on the chromosome. The resulting distance matrices were clustered using Ward’s linkage hierarchical clustering implemented in scipy, and clusters were interpreted as spatial distribution archetypes.

### Statistical Analyses

To test whether LTR-RT abundance, composition, age, and removal patterns differed between centromere architectures, we fitted generalized linear mixed models (GLMMs) with glmmTMB in R (Brooks et al., 2017; R Core Team, 2023; **Supplementary Table S1**). We analysed chromosome-level LTR-RT density (summed annotated LTR-RT length divided by chromosome length), chromosome-level CRM and Ty3-gypsy proportions, element-level LTR identity of intact LTR-RTs, chromosome-level Moran’s I of LTR identity, window- or chromosome-level solo LTR ratios, and Moran’s I of the solo/intact ratio. Chromosome length was included as log10-transformed Mbp and centred for model fitting; model predictions were plotted on the original Mbp scale. For each response we compared nested fixed-effect candidates: null, architecture only, chromosome-size only, additive architecture + size, and architecture × size. Fixed effects were selected with likelihood-ratio tests among nested candidates, and analyses were repeated for the chromosome-size range shared by monocentric and holocentric taxa as a sensitivity analysis.

All final GLMMs retained the same hierarchical taxonomic random-intercept structure (1 | Order/Family/Genus/Species) to approximate phylogenetic non-independence while retaining chromosome- and element-level observations. Element-level LTR identity models additionally included chromosome identity and LTR-RT class as random intercepts; when family-resolved estimated marginal means were calculated, LTR-RT family was instead included as a fixed effect interacting with architecture, and emmeans were evaluated at mean chromosome size. Window-level solo/intact ratio models included chromosome identity as an additional random intercept and were weighted by the number of LTR-RT elements contributing to each ratio. To allow architecture-specific heterogeneity, dispersion was modelled as a function of centromere architecture (dispformula = ∼ arch). Proportional responses were modelled with beta regression after standard boundary adjustments; LTR identity used the length-adjusted transformation ((LTR_identity/100) × (LTR_length - 1) + 0.5) / LTR_length. Moran’s I and log-transformed solo/intact ratios were modelled with Gaussian errors.

We explicitly considered the use of Bayesian phylogenetic frameworks, e.g., *brms* (Bürkner, 2017) or *MCMCglmm* (Hadfield, 2010), which technically support the complex requirements of our study, including multiple observations per species, individual-level data, Beta regression, and nested random effects, while integrating a phylogenetic covariance matrix. However, applying these MCMC methods to a dataset of 1.4 million individual elements was computationally intractable. Furthermore, any data reduction necessary to achieve model convergence, such as aggressive subsampling or aggregating these individual-level observations into species-level means, would have resulted in a substantial loss of biological information, particularly regarding the intra-genomic variance and chromosome size-dependent trends. Therefore, we prioritized the retention of high-resolution, element-level data and utilized hierarchical GLMMs as a robust and comparable alternative for phylogenetic control.

### Clustering of reverse transcriptase sequences and distance-based visualization

To compare lineage structure of dominant LTR-RT families in *Luzula sylvatica* and *Juncus effusus*, we extracted reverse transcriptase protein sequences from full-length element annotations. Sequences were analysed separately for each species and LTR-RT family. Reverse transcriptase sequences were clustered using CD-HIT in protein mode (Li and Godzik, 2006, **Supplementary Table S1**). CD-HIT cluster output files (.clstr) were parsed to summarize the number of clusters, cluster size, and representative centroid sequences at multiple thresholds.

For distance-based visualization, representative reverse transcriptase centroid sequences were selected from the highest identity clustering level with sufficient sequence representation. Selected centroids were aligned using MAFFT with automatic parameter selection (Katoh et al., 2013). Pairwise evolutionary distances between centroid sequences were calculated from the alignment as p-distances, defined as the proportion of mismatched amino acid positions after excluding gap-containing columns. The resulting distance matrix was hierarchically clustered using the average-linkage method and visualized as a heatmap with an accompanying dendrogram. Custom script is available in [https://github.com/437364/nowhere_to_hide_LTR-RTs].

For phylogenetic analysis, reverse transcriptase sequences were grouped by LTR-RT family and aligned using MAFFT (Katoh et al., 2002, 2013) with default parameters. Poorly aligned and gap-rich regions were removed using trimAl (**Supplementary Table S1**) to reduce alignment noise and improve phylogenetic signal. Approximately maximum-likelihood phylogenetic trees were reconstructed for each LTR-RT family using FastTree (**Supplementary Table S1**). Trees were exported in NEWICK format for downstream analyses and visualized using R package ape (**Supplementary Table S1**).

Phylogenetic structure within each LTR-RT family was quantified separately for each species using the Mean Pairwise Distance (MPD) and Mean Nearest Taxon Distance (MNTD) metrics (Webb et al., 2002). MPD captures overall phylogenetic clustering or dispersion among all TE copies within a species, whereas MNTD emphasizes terminal clustering and is more sensitive to recent transposition events. For each family, observed MPD and MNTD values were compared against null distributions generated by random permutation of species labels across tree tips, thereby controlling for family-specific tree topology and sampling effects.

Standardized z-scores were calculated to measure deviation from null expectations, with negative values indicating phylogenetic clustering consistent with recent or ongoing TE expansion, and positive values indicating phylogenetic overdispersion consistent with older or suppressed activity. Families with significant permutation-based differences between species were visualized by comparing species-specific MPD and MNTD z-scores, enabling identification of shared versus lineage-specific TE expansion patterns. Scripts for reverse transcriptase sequence processing, analysis, and visualization are available in the [https://github.com/437364/nowhere_to_hide_LTR-RTs].

### Analysis of ChIP-seq and methyl-seq data

ChIP-seq and Methyl-seq data was sourced from Mata-Sucre et al. 2024 (*L. sylvatica*) and Dias et al. 2024 (*J. effusus*). Analysis workflow followed the methodology in these publications and is available in the [https://github.com/437364/nowhere_to_hide_LTR-RTs].

### Plant material for FISH

The majority of plant material for analyzing the distribution pattern of solo LTRs and full-length LTR-RTs in metaphase chromosomes comes from the cultivation of the Departments or was collected from natural populations (**Supplementary Table S3**). Plants were cultivated under controlled greenhouse conditions.

### DNA extraction and probe preparation

Total DNA was extracted using the NucleoSpin Plant II kit (Macherey-Nagel). TE probes were obtained by polymerase chain reaction (PCR) using specific primers (**Supplementary Table S4**). Primers were designed from available genome assemblies or short-read sequencing (overview for each species is listed in **Supplementary Table S3**). For species with genome assemblies, we first annotated LTR-RTs using the DANTE-LTR tool (Novák et al., 2024). High-confidence full-length elements were clustered by sequence identity using CD-HIT (Li and Godzik, 2006). For each species, we selected two representative elements from the two most abundant LTR-RT families and designed primers targeting both LTR regions and internal GAG or INT regions with Geneious 2024.0.5 (https://www.geneious.com). For species without genome assembly (**Supplementary Table S3**), short reads were obtained from public databases or generated by low-coverage paired-end Illumina sequencing (NovaSeq X platform, 2 × 150 bp) After quality filtering, reads were analysed with the RepeatExplorer pipeline to identify the most abundant LTR-RT family superclusters. Contigs assembled from these superclusters were then used for primer design in Geneious. The code used and additional notes on primer design are available on Github [https://github.com/437364/nowhere_to_hide_LTR-RTs]. PCR-amplified probes were labeled with AF488-dUTP or Cy3-dUTP using the HighFidelity PCR labelling kit.

### Chromosome preparation and fluorescence *in situ* hybridization

For chromosome preparation, we followed a modified protocol by Mandáková and Lysak (2016a). Briefly, root tips from germinated seeds or plants were pre-treated for up to 27 h in ice-cold distilled water and fixed overnight in Carnoy’s fixative (3:1 v/v ethanol : acetic acid). Fixed root tips were stored in 70% ethanol at 4°C until use. After washing in 1×phosphate-buffered saline (1×PBS; pH 7.4), root tips were digested in an enzyme mixture containing 1% (w/v) cellulase R-10 (Duchefa Biochemie), cellulysin cellulase (Merck Millipore), pectolyase Y23 (Duchefa Biochemie), cytohelicase (Sigma-Aldrich), and 20% (v/v) pectinase (Sigma-Aldrich) dissolved in 1×PBS at 37°C. Slides were prepared by spreading meristematic tissue in a drop of 60% acetic acid on a hot plate at 50°C. Afterward, slides were post-fixed in Carnoy’s fixative and 4% formaldehyde solution in 1×PBS, and air-dried.

We performed bicolor fluorescence *in situ* hybridization (FISH) according to a modified protocol by Mandáková and Lysak (2016b). Slides were treated with RNaseA (100 µg/ml) at 37°C for 1 h, and washed in 2×saline-sodium citrate buffer (2×SSC; pH 7.0). Slides were then incubated with pepsin (10 µg/ml) at 37°C for 5 min, washed in 2×SSC, and dehydrated using a graded ethanol series (70, 85, and 100%). A hybridization mixture (50% formamide, 10% dextran sulfate, 2×SSC) combined with labelled probes was denatured onto a slide at 75°C for 5 min and incubated overnight at 37°C in a humid chamber. Post-hybridization washes were performed for 5 min each in 4×SSC at room temperature, 2×SSC at 50°C, and 2×SSC at room temperature. After dehydration, chromosomes were counterstained with 6′-diamidino-2-phenylindole (DAPI; 1.5 μg m/l) in Vectashield (Vector Laboratories).

For slide recycling, we followed a modified protocol by Wakai et al. (2014). Slides were washed in 2×SSC at room temperature for 3 min, treated with 70% formamide dissolved in 2×SSC at 75°C for 5 min, and then washed again in 2×SSC at 75°C for 5 min. After dehydration through ethanol series, the FISH procedure was repeated as described above.

### Microscopy and image analysis

Slides were analyzed with an Olympus BX61 epifluorescence microscope equipped with a ColorView II camera (Olympus) and the Cell^F software (Olympus). Image analysis was performed in ImageJ 1.54f (Schneider et al., 2012; **Supplementary Table S1**). For image analysis, only chromosomes that were clearly separated from neighbouring chromosomes within each metaphase were selected. To identify the boundaries of DAPI-stained chromosomes, the images were enlarged to 300%, and the boundaries were delineated using the ‘Freehand selections’ tool. To evaluate solo-LTR/full-length LTR signals along each chromosome, the ‘Straight line’ was drawn to the chromosome’s central axis, with its width set to cover the entire chromosome. This rectangular selection was transferred to the probe-labelled images and straightened using the ‘Straighten’ function. The grayscale intensities (grey values) of both LTR and GAG/INT probes were measured along the straightened chromosome using the ‘Plot Profile‘ function.

To enable comparison among chromosomes, fluorescence intensities were normalized for each chromosome. Smoothed profiles were calculated using a sliding window average with a window size of 5% of chromosome length. For each window, the ratio of LTR to GAG/INT signal intensity was determined. Because LTR probes hybridize to LTR regions present in solo LTRs as well as in partial and intact LTR-RTs, whereas GAG/INT probes target internal domains expected to be present primarily in intact or internally preserved elements, the LTR to GAG/INT signal ratio was used as a cytogenetic proxy for the relative excess of LTR signal over internal-domain signal. This proxy cannot distinguish solo LTRs from other LTR-containing fragments, but it captures spatial variation in LTR enrichment relative to internal LTR-RT domains along chromosomes.

Spatial autocorrelation of the LTR to GAG/INT signal ratio along chromosomes was quantified with Moran’s I (Moran, 1950) implemented in the ape package (Paradis & Schliep, 2019; **Supplementary Table S1**) in R (R Core Team, 2023). Differences between holocentric and monocentric species were assessed with linear mixed-effects models implemented in the nlme package (Pinheiro et al., 2023; **Supplementary Table S1**), with metaphase nested within slides, species, and families included as nested random effects.

## Results

### 1. Species selection and LTR retrotransposon composition

The total unique holocentric and monocentric species counts included in our study was 40 and 31, respectively. For **bioinformatic analysis**, we selected 37 holocentric and 31 monocentric plant species (**Figure 1a**; **Supplementary Table S2**). In Melanthiaceae, chromosome-level assembly counterpart to holocentric *Chi. japonica* was not available and we used 22 largest sequences from *Cha. luteum* scaffold-level assembly by Kuo et al. (2025). The assembly size ranged from 0.2 (*Cyperus fuscus*) to 17.0 Gbp (*Triticum aestivum*), and the haploid chromosome number (n) varied between 2 (*Rhynchospora tenuis*) and 111 (*Eleocharis dulcis*). The size of chromosome-level scaffolds ranged from 4.4 (*Bolboschoenus planiculmis*) to 868.0 Mbp (*Triticum monococcum*).

In nearly all assemblies BUSCO completeness against the Viridiplantae odb10 dataset ranged between 94–100% with 0 to 3 gaps per Mbp. A notable outlier is *O. longistaminata*, which had 16 gaps per Mbp and was thus excluded from the analysis (**Supplementary Figure S1**). B-chromosomes of *P. chaixii* and *Ca. nigra* were also disregarded. Due to computational constraints, some large Poaceae genomes were represented by only two (*H. vulgare, Triticum* species) or one (*Se. cereale*) chromosome (**Supplementary Table S2**). In total, 1534 chromosome-level scaffolds with detected LTR-RTs were included. Monocentric chromosomes in the dataset were substantially larger than holocentric ones (median 30 Mbp, IQR 19–49 Mbp vs median 10 Mbp, IQR 7–14 Mbp). Because chromosome size could confound architecture-related signals, we included it as a covariate in the generalized linear mixed models.

For **cytogenetic analysis**, we selected 5 holocentric and 4 monocentric species (**Figure 1a**; **Supplementary Table S3**). Cytogenetic analysis was used to expand the number of holocentric species with large chromosomes (*Eleocharis* sp.), which are scarce in the bioinformatics dataset. Some of these species overlap with species selected for bioinformatics analyses to allow for independent verification of the method.

We next examined the composition and abundance of LTR retrotransposons across the analysed species. Because LTR retrotransposon composition is dynamic, substantial variability can be observed even in closely related species. We identified full-length LTR-RTs (defined as elements where both LTRs were identified) and solo LTRs (LTR sequence with no adjacent TE internal regions, see Methods). The **abundance of element families** is **variable** across the taxa, with the *Ale* and *Athila* families dominant in Cyperaceae (e.g., *Carex*, *Cyperus*, *Rhynchospora*), *Angela* in Juncaceae (*Luzula*, *Juncus*), and *Tekay* in Poaceae. (**Fig. 1b, Supplementary Table S5**). **Centromere-associated** elements of the ***CRM* family** were detected in all species examined. Consistent with Neumann et al. (2021), who reported near-complete *CRM* loss in holocentric *Cuscuta*, we found only 2 *CRM* elements in holocentric *Cu. epithymum* versus 198 in monocentric *Cu. campestris*. This reduction was not observed in Cyperaceae or *Chionographis*, indicating *CRM* loss is not a general feature of the transition to holocentricity. In Juncaceae, *Juncus effusus* and both *Luzula* species show a potential reduction of *CRM* abundance (**Supplementary Table S6**). *Galadriel* full-length elements were only found in *Cuscuta* species (22 in *Cu. epithymum* and 3 in *Cu. campestris*) and were excluded from further analysis.

To distinguish the effects of centromere architecture from those of chromosome size and phylogenetic background, we modeled LTR-RT density, Ty3-gypsy proportion, and *CRM* proportion as response variables. Neither overall LTR-RT density nor the relative abundance of Ty3-gypsy elements differed between architectures — in both groups, these increased with chromosome size (**Supplementary Fig. S2a,c,d,f; Supplementary Tables S7, S8**). The exception was *CRM*: monocentric chromosomes had a higher *CRM* proportion among annotated LTR-RTs, and this difference was largest at small to intermediate chromosome sizes, narrowing toward larger chromosomes (**Supplementary Fig. S2b,e**). The architecture-associated signal in LTR-RT composition is therefore specific to *CRM* rather than reflecting a global shift in LTR-RT abundance or Ty3-gypsy content.

The ratio of solo LTRs to full-length elements is likewise specific to the given taxon and LTR-RT family. The four large Poaceae as well as *R. pubera* have a distinctly lower abundance of solo LTRs. In Cyperaceae, *SIRE*, *Tekay*, and *Retand* families show noticeably larger solo LTR ratio, suggesting taxon-specific patterns of ectopic recombination (**Fig. 1b**).

### 2. Age distribution of LTR retrotransposons in monocentric and holocentric species

We used the sequence identity between the 5’ and 3’ LTRs of intact LTR-RTs as a proxy for insertion age. In the raw element-level distribution, holocentric taxa showed higher LTR identity than monocentric taxa (**Fig. 2a**). Species-resolved distributions showed that this pattern was broad but heterogeneous, with the *Athila* family showing a clear contrast between many monocentric and holocentric species (**Fig. 2b**). Similar species-resolved patterns for additional families, including *Angela* and *Ogre*, are shown in **Supplementary Fig. S3**.

**Figure 2.**
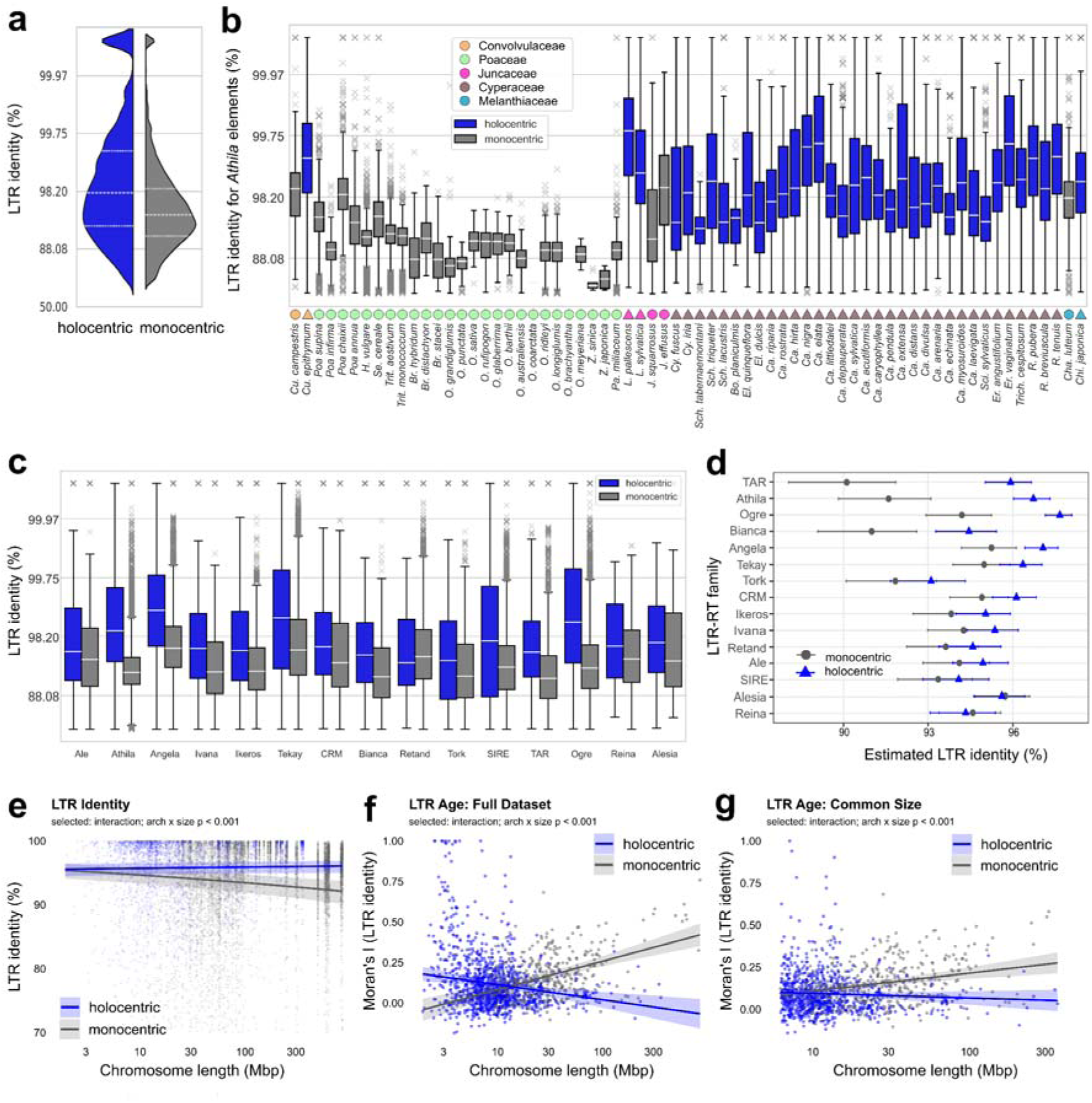
LTR retrotransposons show higher identity and weaker age clustering in holocentric genomes. **(a)** Architecture-level distribution of element-level LTR identity values for intact LTR-RTs in monocentric and holocentric taxa. **(b)** Species-resolved distributions of *Athila* LTR identity. **(c)** Family-level distributions of LTR identity across major LTR-RT families. **(d)** Estimated marginal means of LTR identity for individual LTR-RT families in monocentric and holocentric taxa, derived from a GLMM accounting for chromosome size and hierarchical random effects. Estimates are shown at mean chromosome size and back-transformed from the logit scale; error bars indicate 95% confidence intervals. **(e)** Modelled relationship between LTR identity and chromosome length in the full dataset. **(f)** Modelled relationship between chromosome length and Moran’s I of LTR identity in the full dataset. **(g)** The same Moran’s I analysis restricted to the chromosome-size range shared by monocentric and holocentric taxa. Moran’s I was calculated using a neighbourhood threshold of 0.2, corresponding to 20% of chromosome length. In model-based panels, shaded bands indicate 95% confidence intervals. Grey denotes monocentric taxa and blue denotes holocentric taxa. Additional family- and species-resolved LTR identity distributions, Moran’s I threshold-sensitivity analyses, and Dynamic Time Warping profile analyses are shown in **Supplementary Fig. S3–S5**.

Across major LTR-RT families, holocentric taxa generally showed higher LTR identity than monocentric taxa in the raw distributions or the difference between monocentrics and holocentrics was not significant (**Fig. 2c**). This observation was confirmed by estimated marginal means from the family-level GLMM, which accounted for chromosome size and hierarchical random effects (**Fig. 2d; Supplementary Table S9**).

The element-level mixed model supported an interaction between centromere architecture and chromosome length (**Fig. 2e; Supplementary Tables S7, S8**). In monocentric species, LTR identity decreased with increasing chromosome length, whereas holocentric species maintained high LTR identity across the chromosome-size range. Thus, the model-based analysis supported younger intact LTR-RTs in holocentric genomes, particularly on larger chromosomes.

We next tested whether the spatial clustering of LTR-RT age differed between centromere architectures. Moran’s I of LTR identity was calculated at a neighbourhood threshold of 0.2, corresponding to 20% of chromosome length. In the full dataset, the model selected an interaction between centromere architecture and chromosome length (**Fig. 2f; Supplementary Tables S7, S8**). Moran’s I increased with chromosome length in monocentrics, whereas holocentrics showed a weak or opposite trend. Because monocentric and holocentric taxa partly differed in chromosome-size range, we also analysed the common chromosome-size subset. The interaction remained supported in this restricted dataset, but the negative holocentric slope was attenuated (**Fig. 2g**). Thus, the robust pattern is that larger monocentric chromosomes show stronger spatial clustering of LTR identity, whereas holocentric chromosomes lack this pronounced size-dependent increase.

Threshold-sensitivity analyses of Moran’s I showed the same overall pattern across local to intermediate neighbourhood thresholds, with architecture-related differences weakening at broader thresholds (**Supplementary Figure S4a; Supplementary Table S10a**). A complementary Dynamic Time Warping analysis of chromosome-wide LTR identity profiles showed uniform distribution on holocentric chromosomes, while large monocentric chromosomes often contain younger LTR-RTs on the chromosome arms compared to the central region (**Supplementary Figure S5a**).

### 3. Abundance and distribution of solo LTRs

We next examined solo LTR profiles as signatures of LTR-RT removal. Across species, holocentric taxa tended to show higher solo/intact LTR-RT ratios than monocentric taxa (**Fig. 3a**). Species-resolved distributions showed that this pattern was broad and strongly species-dependent, with particularly low solo/intact ratios in several large-chromosome monocentric grasses and in holocentric *Rhynchospora pubera* (**Fig. 3b**).

**Figure 3.**
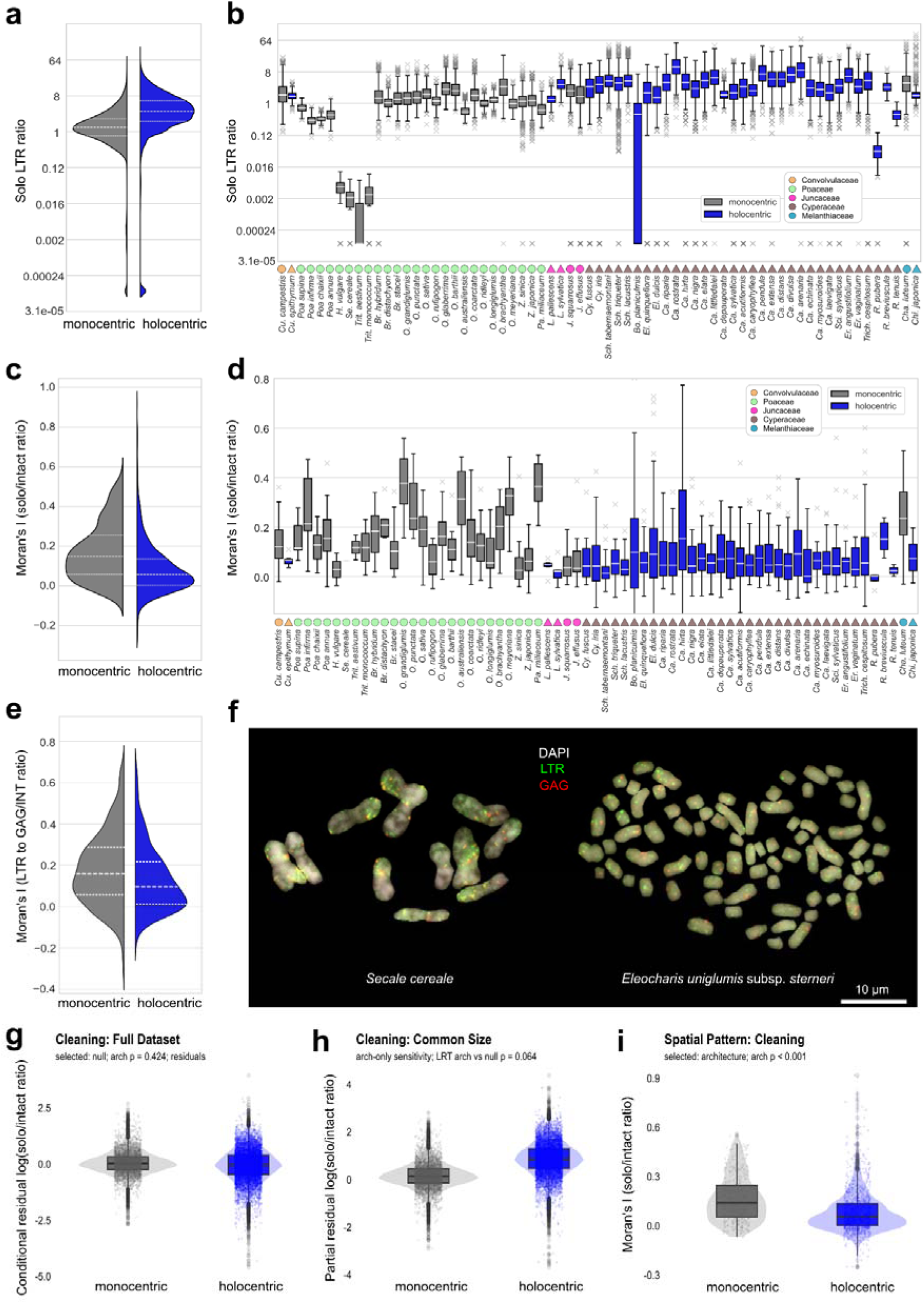
Solo-LTR profiles and spatial removal signatures in monocentric and holocentric chromosomes. **(a)** Distribution of solo/intact LTR-RT ratios across monocentric and holocentric taxa. **(b)** Species-resolved distributions of window-level solo/intact LTR-RT ratios. **(c)** Architecture-level comparison of Moran’s I values calculated from chromosome-level solo/intact LTR-RT ratio profiles. Moran’s I was calculated using a neighbourhood threshold of 0.2, corresponding to 20% of chromosome length. **(d)** Species-resolved distributions of chromosome-level Moran’s I values. Lower Moran’s I indicates weaker spatial clustering of solo/intact ratio along chromosomes. **(e)** Centromere architecture-level comparison of Moran’s I calculated from FISH-based LTR to GAG/INT signal ratios using a neighbourhood threshold of 0.2. **(f)** Representative FISH examples showing LTR and internal-domain signals in monocentric and holocentric chromosomes. **(g)** Mixed-model comparison of overall solo/intact LTR-RT ratio in the full dataset. The selected model contained no fixed effect of centromere architecture, and conditional residuals are shown. **(h)** Sensitivity analysis restricted to the chromosome-size range shared by monocentric and holocentric taxa. The architecture-only model showed a marginal trend toward higher solo/intact ratios in holocentrics. **(i)** Mixed-model comparison of Moran’s I for solo/intact LTR-RT ratio profiles, showing lower spatial autocorrelation in holocentric chromosomes. Colours denote centromere architecture: monocentric taxa in grey and holocentric taxa in blue.

We then quantified the spatial organization of these removal signatures using Moran’s I calculated from the solo/intact LTR-RT ratio. For the main analysis, Moran’s I was calculated with a neighbourhood threshold of 0.2, corresponding to 20% of chromosome length, and the same threshold was used for the genome-based and FISH-based datasets. At this threshold, holocentric chromosomes showed lower Moran’s I values than monocentric chromosomes in the architecture-level comparison (**Fig. 3c**). The species-resolved distributions showed the same pattern, with holocentric species generally exhibiting weaker spatial autocorrelation of solo/intact ratios along chromosomes (**Fig. 3d; Supplementary Table S10b**).

FISH analyses provided an independent cytogenetic view of LTR and internal-domain signal distributions in monocentric and holocentric chromosomes (**Fig. 3e,f; Supplementary Fig. S6; Supplementary Table S11**). Quantitative FISH-based Moran’s I analyses, calculated using the same 0.2 neighbourhood threshold, also showed lower spatial autocorrelation in holocentric chromosomes than in monocentrics (**Supplementary Fig. S7; Supplementary Table S10c,d,e**). Sensitivity analyses across additional Moran’s I thresholds showed that the same direction of effect was recovered across spatial scales, although the difference between architectures became weaker at broader neighbourhood thresholds (**Supplementary Fig. S4c; Supplementary Table S10c**).

A complementary Dynamic Time Warping analysis of chromosome-wide solo/intact ratio profiles recovered a similar spatial contrast. Holocentric chromosomes were predominantly assigned to more uniform removal-profile archetypes, whereas the profile with high removal frequency on chromosome ends was more common on large monocentric chromosomes (**Supplementary Fig. S5b**). Thus, both Moran’s I and time-warp profile clustering indicated weaker spatial structuring of LTR-RT removal signatures in holocentric chromosomes.

To test these patterns while accounting for chromosome size and hierarchical non-independence among observations, we fitted mixed models to the solo-LTR data (**Supplementary Table S7**). In the full dataset, the weighted model of window-level log(solo/intact LTR-RT ratio) selected the null fixed-effect model, and the architecture term was not significant (**Fig. 3g**; p = 0.424). Thus, the model did not support a significant difference in overall solo/intact ratio between monocentric and holocentric taxa in the full dataset. In the common chromosome-size subset, however, the architecture-only sensitivity model showed a marginal trend toward higher solo/intact ratios in holocentrics (**Fig. 3h**; p = 0.064).

In contrast, the model of spatial autocorrelation supported a clear architecture effect. Moran’s I of the solo/intact LTR-RT ratio was significantly lower in holocentric chromosomes than in monocentrics (**Fig. 3i**; p < 0.001), indicating weaker spatial clustering of removal signatures along holocentric chromosomes. The same direction of effect was recovered in the common chromosome-size sensitivity analysis (**Supplementary Fig. S8c**).

### 4. Epigenomic context of LTR retrotransposons in a matched holocentric–monocentric pair

The epigenomic landscapes of monocentric *J. effusus* and holocentric *L. sylvatica* differ substantially, with LTR-RT families associated with distinct epigenetic niches. We analysed the associations with DNA methylation, CenH3 centromere marker, and euchromatin/heterochromatin histone modifications in monocentric *J. effusus* and holocentric *L. sylvatica*. In these species, *Angela*, *Athila*, *Tork*, and *Ivana* are the most prevalent LTR-RT families (**Fig. 4a**).

**Figure 4.**
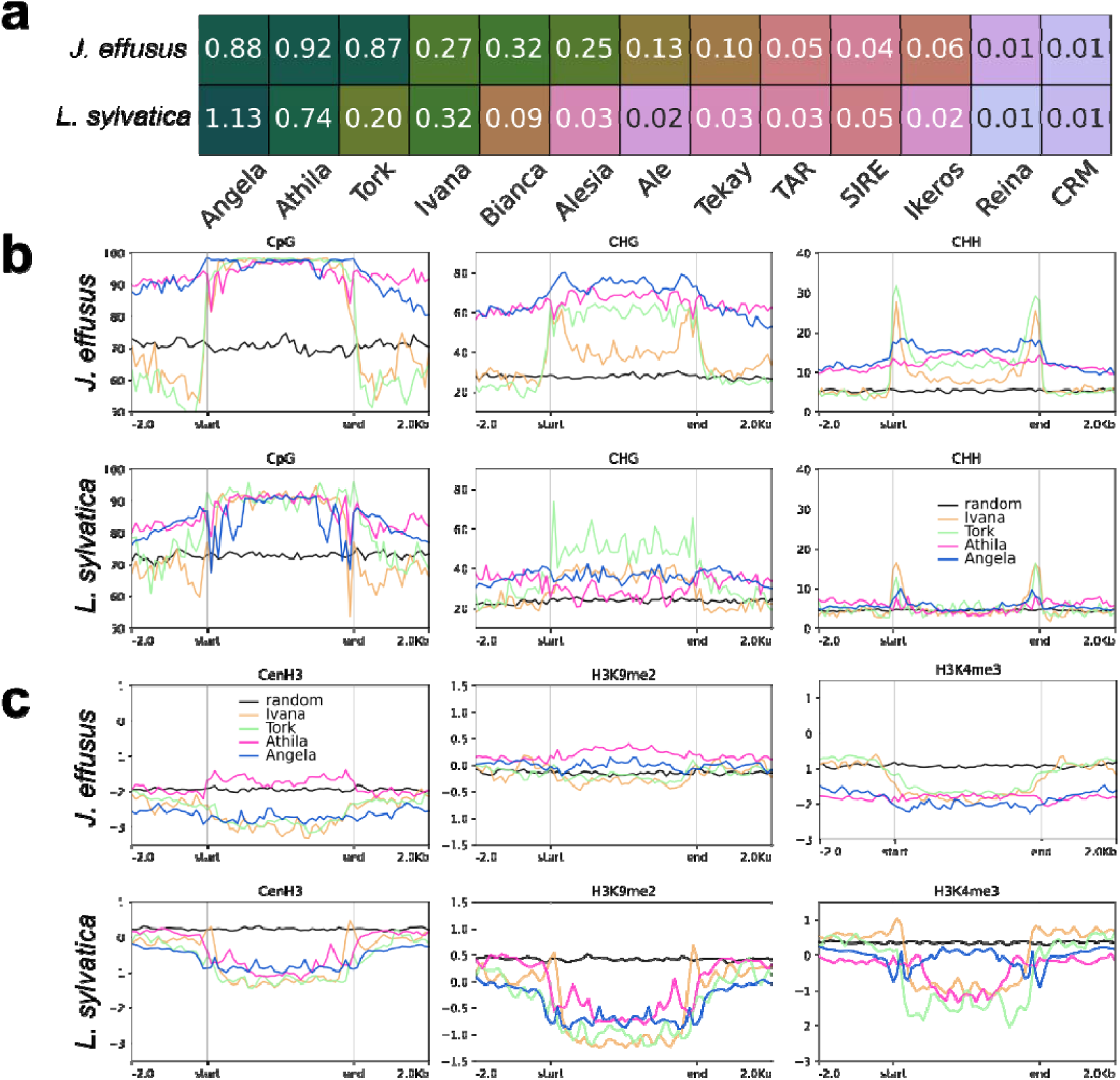
Enrichment of LTR-RTs with epigenetic features in *Luzula sylvatica* and *Juncus effusus*. **(a)** Abundance of LTR-RT families. Labels show the number of intact elements per 1Mbp of assembly. **(b)** Mean methylation profiles of rescaled elements from dominant LTR-RT families and their flanking regions (2kb upstream and downstream; *Angela* - blue, *Athila* - magenta, *Tork* - green, *Ivana* - orange). Individual plots show methylation in CpG, CHG, and CHH contexts. Baseline (black) shows average genomic levels of methylation calculated as an methylation enrichment profile of random genomic regions produced with *bedtools shuffle* command. The Y axis shows the percentage of Cs methylated in a given context. **(c)** Mean CenH3, H3K9me2, and H3K4me3 profiles of rescaled elements from dominant LTR-RT families and their flanking regions (2kb upstream and downstream; *Angela* - blue, *Athila* - magenta, *Tork* - green, *Ivana* - orange). Baseline (black) shows average genomic levels of epigenetic marks calculated as an epigenetic enrichment profile of random genomic regions produced with *bedtools shuffle* command. ChIP-seq signals (Y axis) are shown as log2 (normalized RPKM ChIP/input) and are thus not directly comparable between species.

The *L. sylvatica* epigenomic landscape is dominated by a strong positive correlation between CenH3 and H3K9me2, which is much weaker in *J. effusus* (0.95 vs 0.57). There is also a positive correlation between euchromatin mark H3K4me3 and DNA methylation in all contexts in *L. sylvatica,* whereas the monocentric *J. effusus* displays a negative correlation (∼0.40 vs ∼-0.69, **Supplementary Fig. S9**). As previously reported (Naish & Henderson, 2024; Horáková et al., 2025), CenH3-enriched regions are associated with young LTR-RTs. Consistently, centromeric elements in *J. effusus* and *L. sylvatica* show higher LTR identity than non-centromeric elements (**Supplementary Fig. S10**). In *L. sylvatica*, LTR-RT elements are generally depleted of CenH3, whereas in *J. effusus*, we found a slight CenH3 enrichment of whole *Athila* elements (**Fig. 4b,c**).

Individual LTR-RT families have a distinct epigenetic enrichment profile in each species. *Angela* and *Athila* elements of *J. effusus* are flanked by heterochromatized DNA methylation-rich (H3K4me3 depleted) regions (pericentromeres; Dias et al., 2024), while *Tork* and *Ivana* are flanked by regions with epigenetic enrichment levels near baseline. In *L. sylvatica*, all analyzed element families have similar flanking regions. This further indicates that this heterochromatin-niche preference is absent in holocentric *L. sylvatica*. Solo LTRs show epigenetic profiles similar to LTRs in the intact elements (**Supplementary Fig. S11**).

Reverse transcriptase phylogenies revealed divergent lineage structure between the two species, with *Athila* being the predominant centromeric family in both, yet without evidence of lineage-specific centromeric targeting. (**Fig. 5**). In holocentric *L. sylvatica*, a single large cluster of *Athila* elements dominates the profile, while *J. effusus* displays multiple co-dominant lineages (**Fig. 5a**). This trend is consistent in the *Angela* family (**Supplementary Fig. S12**). We found 108 and 59 full elements overlapping with CenH3-rich regions in *Luzula* and *Juncus*, respectively (**Supplementary Table S12**). *Athila* elements are most frequently inserted into the centromeric regions in both species (∼20% of elements; **Fig. 5c**), even though in *L. sylvatica*, these elements are not themselves enriched for CenH3. When highlighting centromeric elements in the phylogenetic tree (**Fig. 5d**), no specific lineage of either *Athila* or *Angela* appears to be associated with insertion into the centromere. Elements from each species group into distinct subtrees, which reflects interspecies retrotransposon divergence.

**Figure 5.**
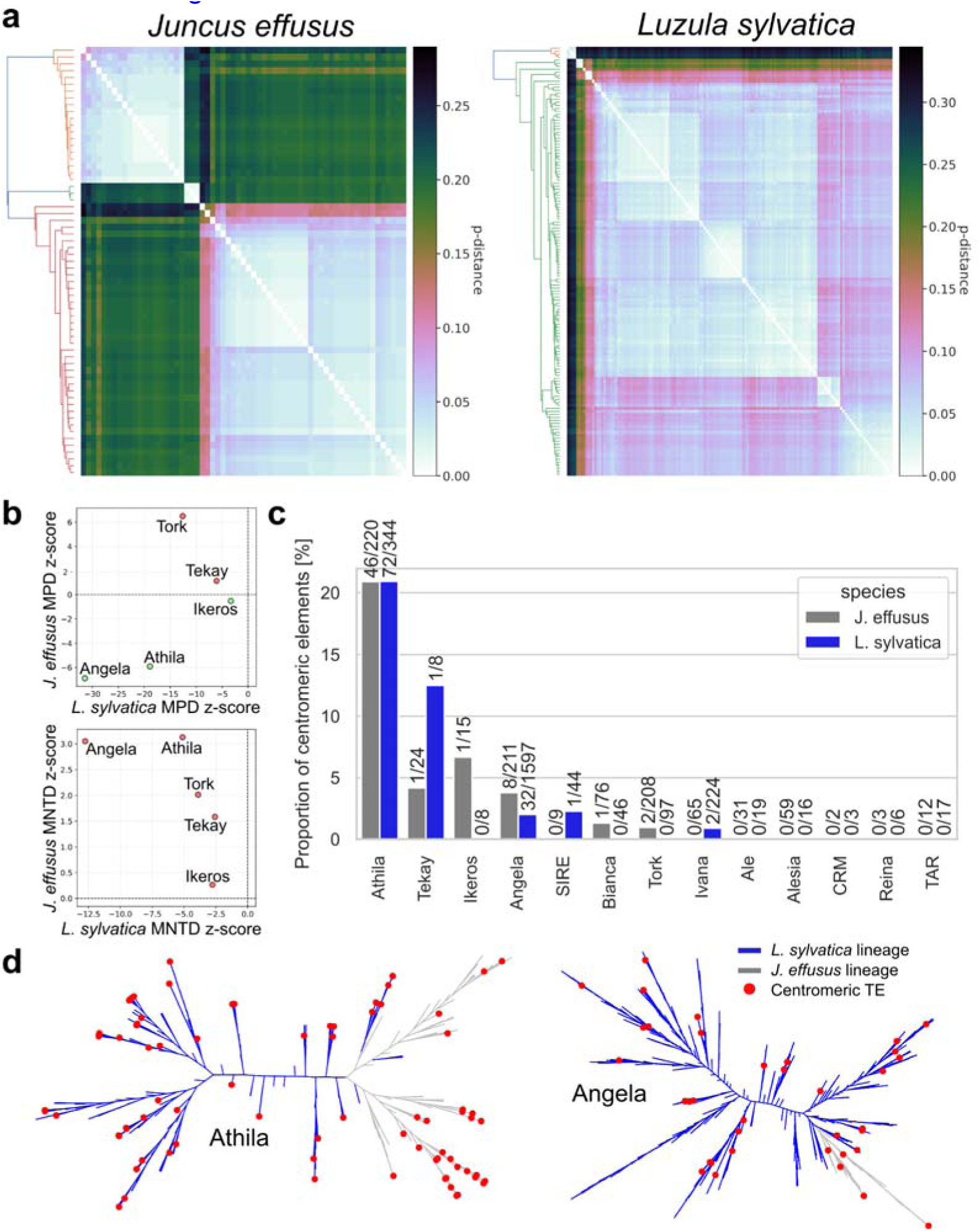
Dominant and centromeric LTR-RT **lineages in *Juncus effusus* and *Luzula sylvatica* (a)** Pairwise distance between clustered reverse transcriptase sequences of *Athila* LTR-RTs **(b)** Comparison of element diversity using mean pairwise distance (MPD) and mean nearest taxon distance (MNTD). **(c)** Proportion centromeric elements. Each bar is labeled with the number of centromeric elements/total number of elements. **(d)** Phylogenetic tree of *Angela* and *Athila* elements from both species. red dots represent elements inserted into the centromeric region.

## Discussion

### 1. LTR retrotransposon family composition reflects lineage history more than centromere architecture

Our results indicate that genome-wide LTR retrotransposon composition and abundance are largely lineage-specific and primarily reflect phylogenetic history rather than centromere organization (**Fig. 1a,b; Supplementary Tables S2, S5, S6**). Cyperaceae, *Luzula*, and *Juncus* were consistently dominated by Ty1-copia elements, whereas grasses were relatively enriched in Ty3-gypsy elements, in agreement with previous reports for these clades (Zedek et al., 2010; Heckmann et al., 2013; Marques et al., 2015; Gebre et al., 2016; Křivánková et al., 2017; Stritt et al., 2020; Mata-Sucre et al., 2024; Yuan et al., 2024; Sader et al., 2026). However, the GLMM analysis showed that Ty3-gypsy proportion increased with chromosome size and did not differ between centromere architectures after chromosome size and hierarchical structure were accounted for (**Supplementary Fig. S2; Supplementary Tables S7, S8**). Similar contrasts in *Chionographis*/*Chamaelirium* and *Cuscuta*, as well as evidence from *Drosera*, further argue against a simple holocentric–monocentric split in Ty1-copia/Ty3-gypsy proportions (this study; Neumann et al., 2021; Kuo et al., 2025; Ávila Robledillo et al., 2025). Thus, the Ty1-copia/Ty3-gypsy balance appears to be shaped mainly by lineage-specific repeatome evolution and chromosome size rather than by centromere architecture (Kumar and Bennetzen, 1999; Terol et al., 2001; Neumann et al., 2011; Pulido and Casacuberta, 2023).

Consistent with this interpretation, overall LTR-RT abundance was not reduced in holocentric species after accounting for chromosome size and phylogenetic structure (**Supplementary Fig. S2; Supplementary Table S7, S8**). Instead, LTR-RT density increased with chromosome size in both monocentric and holocentric taxa, in line with the relationship between chromosome size, recombination density, and repeat accumulation (Bennetzen et al., 2005; Haenel et al., 2018; Brazier and Glémin, 2022; Novák et al., 2020; Zedek et al., 2026). The main exception was CRM: its proportion among annotated LTR-RTs was higher in monocentric chromosomes, especially at small to intermediate chromosome sizes (**Supplementary Fig. S2; Supplementary Tables S7, S8**). Centromere architecture therefore does not determine which LTR-RT lineages dominate a genome, nor does it impose a simple constraint on total LTR-RT abundance. Its strongest effect emerges at another level: the spatial distribution, persistence, and removal of elements.

### 2. Holocentricity reduces pericentromeric refugia for LTR retrotransposons

In monocentric chromosomes, localized centromeres and large heterochromatin blocks, including both pericentromeric and interstitial heterochromatin on chromosome arms, can form broad repeat-rich domains characterized by reduced recombination and epigenetic silencing. Such domains may shelter LTR-RTs from recombination-mediated removal, favouring their retention and local accumulation over evolutionary time (Fransz et al., 2000; Devos et al., 2002; Ma and Bennetzen, 2004; Lamb et al., 2007; Liu et al., 2007; Tian et al., 2009; Evtushenko et al., 2016; Mascher et al., 2017). In holocentric chromosomes, centromere activity is distributed and a single large pericentromeric compartment is absent (Marques et al., 2015; Hofstatter et al., 2022; Castellani et al., 2024; Mata-Sucre et al., 2024). LTR-RTs therefore occupy a more even chromosomal landscape, with fewer opportunities to persist in a protected recombination-poor niche. This is reflected in the weak spatial clustering of LTR-RT age and solo/intact LTR-RT ratios observed here.

The epigenomic contrast between monocentric *Juncus effusus* and holocentric *Luzula sylvatica* provides a candidate mechanistic explanation for this shift, though based on a single matched pair. In *J. effusus*, H3K9me2 correlates with LTR-RTs, satellite repeats, and DNA methylation, consistent with a canonical TE-rich pericentromeric compartment (**Supplementary Fig. S9**; Jackson et al., 2002; Du et al., 2015; Dias et al., 2024). In *L. sylvatica*, H3K9me2 is instead strongly associated with CenH3-marked satellite domains and negatively correlated with LTR-RTs (**Supplementary Fig. S9**; Mata-Sucre et al., 2024). This reassignment of heterochromatin from TE-rich regions to dispersed centromeric units may remove the chromatin-mediated protection that shelters LTR-RTs in monocentric genomes. At the same time, CHH methylation remains associated with LTR-RT boundaries in both species, indicating that RNA-directed DNA methylation still targets elements after the broader heterochromatin landscape has been reorganized (**Fig. 4b,c**; Matzke and Mosher, 2014; Zhang et al., 2018).

### 3. Holocentric genomes show signatures consistent with elevated LTR retrotransposon turnover

If holocentric chromosomes lack broad pericentromeric refugia, LTR-RTs should not only be more uniformly distributed but also more uniformly removed. Solo LTRs arise through unequal homologous recombination between the two LTRs of an element and therefore provide a footprint of recombination-mediated removal (Devos et al., 2002; Ma and Bennetzen, 2004; Vitte and Panaud, 2005; Jedlička et al., 2020). The full-dataset GLMM did not support a significant difference in the overall solo/intact LTR-RT ratio between architectures, although the common chromosome-size subset showed a marginal trend toward higher ratios in holocentrics (**Fig. 3g,h; Supplementary Tables S7, S8**). The more robust signal was spatial: Moran’s I of the solo/intact ratio was lower in holocentric chromosomes, indicating weaker clustering of removal signatures along chromosomes (**Fig. 3c-i; Supplementary Fig. S7, S8**).

Holocentric species also showed higher LTR identity across most LTR-RT families, indicating a younger average element population (**Fig. 2; Supplementary Tables S5, S7-S9**). These age estimates should be interpreted cautiously because LTR identity can be affected by gene conversion and lineage-specific mutation rates (Kijima and Innan, 2010; Cossu et al., 2017; Jedlička et al., 2020; Lewin and Eyre-Walker, 2025). Nevertheless, together with the spatially more uniform removal signatures, the age pattern points to elevated turnover rather than simple repeat depletion: old copies are less likely to persist in protected pericentromeric compartments, while new copies continue to arise through transposition.

The phylogenetic structure of reverse-transcriptase sequences in *L. sylvatica* reinforces this interpretation. Several LTR-RT families showed signatures of recent burst-like amplification, suggesting that the younger age structure of holocentric LTR-RTs reflects both removal of older copies and recent propagation of closely related lineages (**Fig. 5; Supplementary Fig. S12**). This indicates that, at least in this system, holocentricity does not suppress LTR retrotransposon activity *per se*. Rather, it shifts the balance between amplification and removal, leading to a high-turnover regime.

### 4. Chromosome size modulates the architecture effect

LTR-RT density and Ty3-gypsy proportion increased with chromosome size in both architectures, and larger monocentric chromosomes showed stronger spatial clustering of LTR-RT age (**Fig. 2; Supplementary Fig. S2; Supplementary Tables S7, S8**). This may reflect lower recombination density on larger chromosomes, or a shift in recombination outcomes from deletion-producing unequal recombination toward gene conversion, as reported in large plant genomes (Cossu et al., 2017; Haenel et al., 2018; Novák et al., 2020; Zedek et al., 2026). Sensitivity analyses restricted to the chromosome-size range shared by monocentric and holocentric taxa retained the main spatial conclusions but weakened some apparent holocentric trends, especially the solo/intact ratio. The large holocentric genome of *Rhynchospora pubera*, with low solo LTR ratios despite holocentric organization, may represent an important example of this attenuation (**Fig. 3b**).

Together, our results argue against a simple model in which holocentricity determines LTR-RT content. Instead, centromere architecture shapes the chromosomal ecology of LTR retrotransposons. LTR-RT composition largely reflects lineage history, total abundance, and Ty3-gypsy proportion scale with chromosome size, and *CRM* represents the clearest architecture-associated compositional exception. Holocentricity primarily affects the spatial distribution, persistence, and removal of elements. This framework helps reconcile apparently contrasting observations in holocentric plants: they may be TE-rich or TE-poor, may contain recent bursts of particular LTR-RT families, and may nevertheless show young element populations and weak spatial clustering of removal signatures (Zedek et al., 2010; de Souza et al., 2018; Mata-Sucre et al., 2024). These features reflect a dynamic equilibrium in which transposition continues, but long-term persistence in protected pericentromeric compartments is reduced.

## DATA AVAILABILITY

Element table used for linear models analysis and annotations of full, partial, and solo elements are available in Zenodo repository (10.5281/zenodo.21345425). Raw Illumina sequencing data is available in SRA under accession PRJNA1494884.

## CODE AVAILABILITY

Code used in this manuscript is available in Github code repository https://github.com/437364/nowhere_to_hide_LTR-RTs (10.5281/zenodo.21346479).

## AUTHORS’ CONTRIBUTIONS

FZ conceived of the study. MK, KP, PJ, PB, ZK, EK, and FZ designed the study. MK and PJ performed bioinformatic analyses. KP and JŠ performed wet lab work. KP performed cytogenetic and image analyses. MK, KP, and FZ performed statistical analyses. MK, KP, PJ, PB, ZK, AM, EK, and FZ wrote the manuscript. All authors discussed the results and approved the final version of the manuscript.

## ACKNOWLEDGEMENTS

We thank the staff from The Botanical Garden of the Faculty of Science, Masaryk University, for providing seeds. Computational resources were provided by Metacentrum, the e-INFRA CZ project (ID:90254), supported by the Ministry of Education, Youth and Sports of the Czech Republic.

## REFERENCES

Ávila Robledillo L, Fleck SJ, Kirshner J, Becker D, Bhatia A, Bringmann G et al. 2025. Centromere divergence and allopolyploidy reshape carnivorous sundew genomes. preprint, doi: 10.1101/2025.07.27.666937.

Bennetzen JL, Ma J, Devos KM. 2005. Mechanisms of recent genome size variation in flowering plants. Annals of Botany 95(1): 127–132. doi: 10.1093/aob/mci008.

Biémont C, Vieira C. 2006. Junk as an evolutionary force. Nature 443: 521–524. doi: 10.1038/443521a.

Brazier T, Glémin S. 2022. Diversity and determinants of recombination landscapes in flowering plants. PLoS Genetics 18: e1010141. doi: 10.1371/journal.pgen.1010141.

Brooks ME, Kristensen K, van Benthem KJ, Magnusson A, Berg CW, Nielsen A, Skaug HJ, Maechler M, Bolker BM. 2017. glmmTMB Balances Speed and Flexibility Among Packages for Zero-inflated Generalized Linear Mixed Modeling. The R Journal 9(2): 378–400. doi:10.32614/RJ-2017-066.

Bureš P, Elliott TL, Veselý P, Šmarda P, Forest F, Leitch IJ, Nic Lughadha E, Soto Gomez M, Pironon S, Brown MJM, Šmerda J, Zedek F. 2024. The global distribution of angiosperm genome size is shaped by climate. The New Phytologist 242(2): 744–759. doi: 10.1111/nph.19544.

Bureš P, Zedek F. 2014. Holokinetic drive: centromere drive in chromosomes without centromeres. Evolution 68(8): 2412–2420. doi: 10.1111/evo.12437.

Bureš P, Zedek F, Marková M. 2013. Holocentric chromosomes. In: Leitch IJ, Greilhuber J, Doležel J, Wendel JF (eds). Plant Genome Diversity Volume 2 Physical Structure, Behaviour and Evolution of Plant Genomes. Springer Verlag Wien, p. 187–208. doi: 10.1007/978-3-7091-1160-4_12.

Bürkner P. 2017. brms: An R Package for Bayesian Multilevel Models Using Stan. Journal of Statistical Software 80: 1–28. doi:10.18637/jss.v080.i01.

Castellani M, Zhang M, Thangavel G, Mata-Sucre Y, Lux T, Campoy JA, Marek M, Huettel B, Sun H, Mayer KFX, Schneeberger K, Marques A. 2024. Meiotic recombination dynamics in plants with repeat-based holocentromeres shed light on the primary drivers of crossover patterning. Nature Plants 10: 423–438. doi: 10.1038/s41477-024-01625-y.

Charles M, Belcram H, Just J, Huneau C, Viollet A, Couloux A, Segurens B, Carter M, Huteau V, Coriton O, Appels R, Samain S, Chalhoub B. 2008. Dynamics and differential proliferation of transposable elements during the evolution of the B and A genomes of wheat. Genetics 180(2): 1071–1086. doi: 10.1534/genetics.108.092304.

Cossu RM, Casola C, Giacomello S, Vidalis A, Scofield DG, Zuccolo A. 2017. LTR Retrotransposons Show Low Levels of Unequal Recombination and High Rates of Intraelement Gene Conversion in Large Plant Genomes. Genome Biology and Evolution 9(12): 3449–3462. doi: 10.1093/gbe/evx260.

de Souza TB, Chaluvadi SR, Johnen L, Marques A, González-Elizondo MS, Bennetzen JL, Vanzela ALL. 2018. Analysis of retrotransposon abundance, diversity and distribution in holocentric *Eleocharis* (Cyperaceae) genomes. Annals of Botany 122(2): 279–290. doi: 10.1093/aob/mcy066.

Deniz Ö, Frost JM, Branco MR. 2019. Regulation of transposable elements by DNA modifications. Nature Reviews Genetics 20(7): 417–431. doi: 10.1038/s41576-019-0106-6.

Devos KM, Brown JK, Bennetzen JL. 2002. Genome size reduction through illegitimate recombination counteracts genome expansion in *Arabidopsis*. Genome Research 12(7): 1075–1079. doi: 10.1101/gr.132102.

Dias Y, Mata-Sucre Y, Thangavel G, Costa L, Báez M, Houben A, Marques A, Pedrosa-Harand A. 2024. How diverse a monocentric chromosome can be? Repeatome and centromeric organization of *Juncus effusus* (Juncaceae). The Plant Journal 118: 1832–1847. doi: 10.1111/tpj.16712.

Du J, Tian Z, Hans CS, Laten HM, Cannon SB, Jackson SA, Shoemaker RC, Ma J. 2010. Evolutionary conservation, diversity and specificity of LTR-retrotransposons in flowering plants: insights from genome-wide analysis and multi-specific comparison. The Plant Journal : for cell and molecular biology 63(4): 584–598. doi: 10.1111/j.1365-313X.2010.04263.x.

Du J, Johnson LM, Jacobsen SE, Patel DJ. 2015. DNA methylation pathways and their crosstalk with histone methylation. Nat Rev Mol Cell Biol 16(9): 519–532. doi: 10.1038/nrm4043.

Evtushenko EV, Levitsky VG, Elisafenko EA, Gunbin KV, Belousov AI, Šafář J, Doležel J, Vershinin AV. 2016. The expansion of heterochromatin blocks in rye reflects the co-amplification of tandem repeats and adjacent transposable elements. BMC Genomics 17: 337.doi: 10.1186/s12864-016-2667-5.

Fransz PF, Armstrong S, de Jong JH, Parnell LD, van Drunen C, Dean C, Zabel P, Bisseling T, Joneso GH. 2000. Integrated cytogenetic map of chromosome arm 4S of *Arabidopsis thaliana*: structural organization of heterochromatic knob and centromere region. Cell 100(3): 367–376. doi: 10.1016/S0092-8674(00)80672-8.

Forest F. 2023. Species-level phylogenetic trees of all angiosperm species (100 trees) [Data set]. Zenodo. 10.5281/zenodo.7600341.

Gebre YG, Bertolini E, Pè ME, Zuccolo A. 2016. Identification and characterization of abundant repetitive sequences in *Eragrostis tef* cv. Enatite genome. BMC Plant Biology 16: 39. doi: 10.1186/s12870-016-0725-4.

Hadfield JD. 2010. MCMC Methods for Multi-Response Generalized Linear Mixed Models: The MCMCglmm R Package. Journal of Statistical Software 33: 1–22. https://www.jstatsoft.org/v33/i02/.

Haenel Q, Laurentino TG, Roesti M, Berner D. 2018. Meta-analysis of chromosome-scale crossover rate variation in eukaryotes and its significance to evolutionary genomics. Molecular Ecology 27: 2477–2497. doi: 10.1111/mec.14699.

Heckmann S, Macas J, Kumke K, Fuchs J, Schubert V, Ma L, Novák P, Neumann P, Taudien S, Platzer M, Houben A. 2013. The holocentric species *Luzula elegans* shows interplay between centromere and large-scale genome organization. The Plant Journal : for Cell and Molecular Biology 73(4): 555–565. doi: 10.1111/tpj.12054.

Hobza R, Cegan R, Jesionek W, Kejnovsky E, Vyskot B, Kubat Z. 2017. Impact of Repetitive Elements on the Y Chromosome Formation in Plants. Genes 8(11): 302. doi: 10.3390/genes8110302.

Hofstatter PG, Thangavel G, Castellani M, Marques A. 2021. Meiosis Progression and Recombination in Holocentric Plants: What Is Known? Frontiers in Plant Science 12: 658296. doi: 10.3389/fpls.2021.658296.

Hofstatter PG, Thangavel G, Lux T, Neumann P, Vondrak T, Novak P, et al. 2022. Repeat-based holocentromeres influence genome architecture and karyotype evolution. Cell 185(17): 3153–3168.e18. doi: 10.1016/j.cell.2022.06.045.

Horáková L, Jedlička P, Čegan R, Navrátilová P, Tanaka H, Toyoda A, Itoh T, Akagi T, Ono E, Hudzieczek V, Patzak J, Šafář J, Hobza R, Bačovský V. 2025. Dynamic patterns of repeats and retrotransposons in the centromeres of *Humulus lupulus* L. The New Phytologist 247:2766–2780. doi: 10.1111/nph.70380.

Horváth V, Merenciano M, González J. 2017. Revisiting the Relationship between Transposable Elements and the Eukaryotic Stress Response. Trends in Genetics 33(11): 832–841. doi: 10.1016/j.tig.2017.08.007.

Jackson JP, Lindroth AM, Cao X, Jacobsen SE. 2002. Control of CpNpG DNA methylation by the KRYPTONITE histone H3 methyltransferase. Nature 416: 556–560. doi: 10.1038/nature731.

Jedlicka P, Lexa M, Kejnovsky E. 2020. What Can Long Terminal Repeats Tell Us About the Age of LTR Retrotransposons, Gene Conversion and Ectopic Recombination? Frontiers in Plant Science 11: 644. doi: 10.3389/fpls.2020.00644.

Katoh K, Misawa K, Kuma K, Miyata T. 2002. MAFFT: a novel method for rapid multiple sequence alignment based on fast Fourier transform. Nucleic Acids Res 30(14): 3059–3066. doi: 10.1093/nar/gkf436.

Katoh K, Standley DM. 2013. MAFFT multiple sequence alignment software version 7: improvements in performance and usability. Mol Biol Evol 30(4): 772–780. doi: 10.1093/molbev/mst010.

Kejnovsky E, Hobza R, Kubat Z, Cermak T, Vyskot B. 2009. The role of repetitive DNA in structure and evolution of sex chromosomes in plants. Heredity 102: 533–541. doi: 10.1038/hdy.2009.17.

Kejnovsky E, Hawkins JS, Feschotte C. 2012. Plant Transposable Elements: Biology and Evolution. In: Wendel J, Greilhuber J, Dolezel J, Leitch I (eds). Plant Genome Diversity Volume 1. Springer, Vienna, p. 17–34. doi: 10.1007/978-3-7091-1130-7_2.

Kejnovsky E, Jedlicka P. 2022. Nucleic acids movement and its relation to genome dynamics of repetitive DNA. BioEssays 44(4): 2100242. doi: 10.1002/bies.202100242.

Kejnovsky E, Jedlicka P, Lexa M, Kubat Z. 2025. Factors determining chromosomal localization of transposable elements in plants. Plant Biology 27(6): 975–989. doi: 10.1111/plb.70057.

Kent TV, Uzunović J, Wright SI. 2017. Coevolution between transposable elements and recombination. Philosophical transactions of the Royal Society of London. Series B, Biological sciences 372(1736): 20160458. doi: 10.1098/rstb.2016.0458.

Kijima TE, Innan H. 2010. On the estimation of the insertion time of LTR retrotransposable elements. Molecular Biology and Evolution 27(4): 896–904. doi: 10.1093/molbev/msp295.

Křivánková A, Kopecký D, Stočes Š, Doležel J, Hřibová E. 2017. Repetitive DNA: a versatile tool for karyotyping in *Festuca pratensis* huds. Cytogenetic and Genome Research 151(2): 96–105. doi: 10.1159/000462915.

Kumar A, Bennetzen JL. 1999. Plant retrotransposons. Annual Review of Genetics 33: 479–532. doi: 10.1146/annurev.genet.33.1.479.

Kuo YT, Neumann P, Chen J, Fuchs J, Schubert V, Kumke K, Sader MA, Melzer M, Zhu Z, Himmelbach A, Hentrich H, Macas J, Houben A. 2025. Kinetochore mutations and histone phosphorylation pattern changes accompany holo- and macro-monocentromere evolution. Nature Communications 16(1): 11332. doi: 10.1038/s41467-025-67524-8.

Lamb JC, Yu W, Han F, Birchler JA. 2007. Plant chromosomes from end to end: telomeres, heterochromatin and centromeres. Current Opinion in Plant Biology 10(2): 116–122. doi: 10.1016/j.pbi.2007.01.008.

Lewin L, Eyre-Walker A. 2025. Estimates of the Mutation Rate per Year Can Explain Why the Molecular Clock Depends on Generation Time. Molecular Biology and Evolution 42(4): msaf069. doi: 10.1093/molbev/msaf069.

Li W, Godzik A. 2006. Cd-hit: a fast program for clustering and comparing large sets of protein or nucleotide sequences. Bioinformatics 22(13): 1658–1659. doi: 10.1093/bioinformatics/btl158.

Lichten M, Borts RH, Haber JE. 1987. Meiotic gene conversion and crossing over between dispersed homologous sequences occurs frequently in *Saccharomyces cerevisiae*. Genetics 115(2): 233–246. doi: 10.1093/genetics/115.2.233.

Liu R, Vitte C, Ma J, Mahama AA, Dhliwayo T, Lee M, Bennetzen JL. 2007. A GeneTrek analysis of the maize genome. Proceedings of the National Academy of Sciences 104(29): 11844–11849. doi: 10.1073/pnas.0704258104.

Lynch M, Bobay LM, Catania F, Gout JF, Rho M. 2011. The repatterning of eukaryotic genomes by random genetic drift. Annual Review in Genomics and Human Genetics 12: 347–366. doi: 10.1146/annurev-genom-082410-101412.

Ma J, Bennetzen JL. 2004. Rapid recent growth and divergence of rice nuclear genomes. Proc Natl Acad Sci USA 101(34): 12404–12410. doi: 10.1073/pnas.0403715101.

Makarevitch I, Waters AJ, West PT, Stitzer MC, Hirsch CN, Ross-Ibarra J, Springer NM. 2015. Transposable elements contribute to activation of maize genes in response to abiotic stress. PLoS Genetics 11(1): e1004915. doi: 10.1371/journal.pgen.1004915.

Mandáková T, Lysak MA. 2016a. Chromosome preparation for cytogenetic analyses in *Arabidopsis*. Current Protocols in Plant Biology 1: 43–51. doi: 10.1002/cppb.20009.

Mandáková T, Lysak MA. 2016b. Painting of *Arabidopsis* chromosomes with chromosome-specific BAC Clones. Current Protocols in Plant Biology 1: 359–371. doi: 10.1002/cppb.20022.

Mandrioli M, Manicardi GC. 2012. Unlocking holocentric chromosomes: new perspectives from comparative and functional genomics? Current Genomics 13: 343–349. doi: 10.2174/138920212801619250.

Mandrioli M, Manicardi GC. 2020. Holocentric chromosomes. PLoS Genetics 16(7): e1008918. doi: 10.1371/journal.pgen.1008918.

Marques A, Drinnenberg IA. 2025. Same but different: Centromere regulations in holocentric insects and plants. Curr Opin Cell Biol 93:102484. doi: 10.1016/j.ceb.2025.102484.

Marques A, Ribeiro T, Neumann P, Macas J, Novák P, Schubert V, Pellino M, Fuchs J, Ma W, Kuhlmann M, Brandt R, Vanzela ALL, Beseda T, Šimková H, Pedrosa-Harand A, Houben A. 2015. Holocentromeres in *Rhynchospora* are associated with genome-wide centromere-specific repeat arrays interspersed among euchromatin. Proceedings of the National Academy of Sciences 112: 13633–13638. doi: 10.1073/pnas.1512255112.

Mascher M, Gundlach H, Himmelbach A, Beier S, Twardziok SO, Wicker T, Radchuk V, Dockter C, Hedley PE, Russell J, Bayer M, Ramsay L, Liu H, Haberer G, Zhang XQ, Zhang Q, Barrero RA, Li L, Taudien S, Groth M, Felder M, Hastie A, Šimková H, Staňková H, Vrána J, Chan S, Muñoz-Amatriaín M, Ounit R, Wanamaker S, Bolser D, Colmsee C, Schmutzer T, Aliyeva-Schnorr L, Grasso S, Tanskanen J, Chailyan A, Sampath D, Heavens D, Clissold L, Cao S, Chapman B, Dai F, Han Y, Li H, Li X, Lin C, McCooke JK, Tan C, Wang P, Wang S, Yin S, Zhou G, Poland JA, Bellgard MI, Borisjuk L, Houben A, Doležel J, Ayling S, Lonardi S, Kersey P, Langridge P, Muehlbauer GJ, Clark MD, Caccamo M, Schulman AH, Mayer KFX, Platzer M, Close TJ, Scholz U, Hansson M, Zhang G, Braumann I, Spannagl M, Li C, Waugh R, Stein N. 2017. A chromosome conformation capture ordered sequence of the barley genome. Nature 544(7651): 427–433. doi: 10.1038/nature22043.

Mata-Sucre Y, Krátká M, Oliveira L, Neumann P, Macas J, Schubert V, Huettel B, Kejnovský E, Houben A, Pedrosa-Harand A, Souza G, Marques A. 2024. Repeat-based holocentromeres of the woodrush *Luzula sylvatica* reveal insights into the evolutionary transition to holocentricity. Nature Communications 15, 9565. doi: 10.1038/s41467-024-53944-5.

McVean. 2010. What drives recombination hotspots to repeat DNA in humans? *Philosophical transactions of the Royal Society of London. Series B*, Biological sciences 365(1544): 1213–1218. doi: 10.1098/rstb.2009.0299.

Morales-Díaz N, Sushko S, Campos-Dominguez L, Kopalli V, Golicz AA, Castanera R, Casacuberta JM. 2025. Tandem LTR-retrotransposon structures are common and highly polymorphic in plant genomes. Mobile DNA 16(1): 10. doi: 10.1186/s13100-025-00347-y.

Moran PAP. 1950. Notes on Continuous Stochastic Phenomena. Biometrika 37: 17–23. doi: 10.2307/2332142.

Myers S, Freeman C, Auton A, Donnelly P, McVean G. 2008. A common sequence motif associated with recombination hot spots and genome instability in humans. Nature Genetics 40(9): 1124–1129. doi: 10.1038/ng.213.

Naish M, Henderson IR. 2024. The structure, function, and evolution of plant centromeres. Genome Research 34:161–178. doi: 10.1101/gr.278409.123.

Neumann P, Navrátilová A, Koblížková A, Kejnovský E, Hřibová E, Hobza R, Widmer A, Doležel J, Macas J. 2011. Plant centromeric retrotransposons: a structural and cytogenetic perspective. Mobile DNA 2(1): 4. doi: 10.1186/1759-8753-2-4.

Neumann P, Oliveira L, Čížková J, Jang TS, Klemme S, Novák P, Stelmach K, Koblížková A, Doležel J, Macas J. 2021. Impact of parasitic lifestyle and different types of centromere organization on chromosome and genome evolution in the plant genus *Cuscuta*. The New Phytologist 229(4): 2365–2377. doi: 10.1111/nph.17003.

Novak P, Guignard MS, Neumann P, Kelly LJ, Mlinarec J, Koblížková A, Dodsworth S, Kovařík A, Pellicer J, Wang W, Macas J, Leitch IJ, Leitch AR. 2020. Repeat-sequence turnover shifts fundamentally in species with large genomes. Nature Plants 6(11): 1325–1329. doi: 10.1038/s41477-020-00785-x.

Novák P, Hoštáková N, Neumann P, Macas J. 2024. DANTE and DANTE_LTR: lineage-centric annotation pipelines for long terminal repeat retrotransposons in plant genomes. NAR Genom Bioinform. 6:lqae113. doi: 10.1093/nargab/lqae113.

Oliver KR, Greene WK. 2009. Transposable elements: powerful facilitators of evolution. Bioessays 31(7): 703–714. doi: 10.1002/bies.200800219.

Ou S, Jiang N. 2018. LTR_retriever: A Highly Accurate and Sensitive Program for Identification of Long Terminal Repeat Retrotransposons. Plant Physiol 176(2):1410–1422. doi: 10.1104/pp.17.01310.

Paradis E, Schliep K. 2019. ape 5.0: an environment for modern phylogenetics and evolutionary analyses in R. Bioinformatics 35, 526–528. doi: 10.1093/bioinformatics/bty633.

Pinheiro J, Bates D, R Core Team. 2023. nlme: Linear and Nonlinear Mixed Effects Models. R package version 3.1–162. https://CRAN.R-project.org/package=nlme.

Pulido M, Casacuberta JM. 2023. Transposable element evolution in plant genome ecosystems. Current Opinion in Plant Biology 75: 102418. doi: 10.1016/j.pbi.2023.102418.

R Core Team. 2023. R: a language and environment for statistical computing. Vienna, Austria: R Foundation for Statistical Computing. http://www.Rproject.org/.

Rebollo R, Horard B, Hubert B, Vieira C. 2010. Jumping genes and epigenetics: Towards new species. Gene 454(1–2): 1–7. doi: 10.1016/j.gene.2010.01.003.

Robinson O, Dylus D, Dessimoz C. 2016. Phylo.io: Interactive Viewing and Comparison of Large Phylogenetic Trees on the Web. Molecular Biology and Evolution 33(8): 2163–2166. doi: 10.1093/molbev/msw080.

Sader M, Mata-Sucre Y, Kuo Y-T, Schubert V, Nascimento T, Fuchs J, Dias Y, Pistrick K, Sargheini N, Huettel B, Vanzela ALL, Marques A, Houben A, Pedrosa-Harand A. 2026. Holocentromere diversity in Cyperaceae: contrasting repeat organisation in Mapanioideae and Cyperoideae. preprint, doi: 10.64898/2026.06.15.732478.

Schnable PS, Ware D, Fulton RS, Stein JC, Wei F, Pasternak S, Liang C, Zhang J, et al. 2009. The B73 maize genome: complexity, diversity, and dynamics. Science 326(5956): 1112–1115. doi: 10.1126/science.1178534.

Schneider CA, Rasband WS, Eliceiri KW. 2012. NIH Image to ImageJ: 25 years of image 985 analysis. Nature Methods 9: 671–675. doi: 10.1038/nmeth.2089.

Seppey M, Manni M, Zdobnov EM. 2019. BUSCO: Assessing Genome Assembly and Annotation Completeness. Methods in Molecular Biology 1962: 227–245. doi: 10.1007/978-1-4939-9173-0_14.

Shipilina D, Näsvall K, Höök L, Vila R, Talavera G, Backström N. 2022. Linkage mapping and genome annotation give novel insights into gene family expansions and regional recombination rate variation in the painted lady (*Vanessa cardui*) butterfly. Genomics 114: 110481. doi: 10.1016/j.ygeno.2022.110481.

Simon L, Voisin M, Tatout C, Probst AV. 2015. Structure and Function of Centromeric and Pericentromeric Heterochromatin in *Arabidopsis thaliana*. Frontiers in Plant Science 6: 1049. doi: 10.3389/fpls.2015.01049.

Smit AFA, Hubley R, Green P. 2013–2015. RepeatMasker Open-4.0. http://www.repeatmasker.org.

Stritt C, Wyler M, Gimmi EL, Pippel M, Roulin AC. 2020. Diversity, dynamics and effects of long terminal repeat retrotransposons in the model grass *Brachypodium distachyon*. The New Phytologist 227(6): 1736–1748. doi: 10.1111/nph.16308.

Terol J, Castillo MC, Bargues M, Pérez-Alonso M, de Frutos R. 2001. Structural and evolutionary analysis of the copia-like elements in the *Arabidopsis thaliana* genome. Molecular Biology and Evolution 18(5): 882–892. doi: 10.1093/oxfordjournals.molbev.a003870.

Tian Z, Rizzon C, Du J, Zhu L, Bennetzen JL, Jackson SA, Gaut BS, Ma J. 2009. Do genetic recombination and gene density shape the pattern of DNA elimination in rice long terminal repeat retrotransposons? Genome Research 19: 2221–2230. doi: 10.1101/gr.083899.108.

Vitte C, Panaud O. 2005. LTR retrotransposons and flowering plant genome size: emergence of the increase/decrease model. Cytogenetic and Genome Research 110(1-4): 91–107. doi: 10.1159/000084941.

Wakai S, Shibuki Y, Yokozawa K, Nakamura S, Adegawa Y, Yoshida A, Tsuta K, Furuta K. 2014. Recycling and Long-Term Storage of *Fluorescence In Situ Hybridization* Slides. American Journal of Clinical Pathology 141(3): 374–380. doi: 10.1309/AJCPYX1UTI7LDAUY.

Wang Z, Zhang J, Yang W, An N, Zhang P, Zhang G, Zhou Q. 2014. Temporal genomic evolution of bird sex chromosomes. BMC Evolutionary Biology 14: 250. doi: 10.1186/s12862-014-0250-8.

Wang C, Chen Z, Copenhaver GP, Wang Y. 2024. Heterochromatin in plant meiosis. Nucleus 15(1): 2328719. doi: 10.1080/19491034.2024.2328719.

Webb C, Ackerly D, McPeek M, Donoghue M. 2002. Phylogenies and community ecology. Annual Review of Ecology and Systematics 33: 475–505. doi: 10.1146/annurev.ecolsys.33.010802.150448.

Wells JN, Feschotte C. 2020. A field guide to eukaryotic transposable elements. Annual Review of Genetics 54: 539–561. doi: 10.1146/annurev-genet-040620-022145.

Wicker T, Sabot F, Hua-Van A, Bennetzen JL, Capy P, Chalhoub B, Flavell A, Leroy P, Morgante M, Panaud O, Paux E, SanMiguel P, Schulman AH. 2007. A unified classification system for eukaryotic transposable elements. Nature Reviews Genetics 8(12): 973–982. doi: 10.1038/nrg2165.

Wicker T, Gundlach H, Spannagl M, Uauy C, Borrill P, Ramírez-González RH, De Oliveira R; International Wheat Genome Sequencing Consortium; Mayer KFX, Paux E, Choulet F. 2018. Impact of transposable elements on genome structure and evolution in bread wheat. Genome Biology 19(1): 103. doi: 10.1186/s13059-018-1479-0.

Yuan T, Gao X, Xiang N, Wei P, Zhang G. 2024. The genome assembly of *Carex breviculmis* provides evidence for its phylogenetic localization and environmental adaptation. Annals of Botany 134(3): 467–484. doi: 10.1093/aob/mcae085.

Zedek F, Šmerda J, Šmarda P, Bureš P. 2010. Correlated evolution of LTR retrotransposons and genome size in the genus *Eleocharis*. BMC Plant Biology 10: 265. doi: 10.1186/1471-2229-10-265.

Zedek F, Bureš P, Elliott TL, Escudero M, Lucek K, Marques A. 2026. Chromosome size as a robust predictor of recombination rate: insights from holocentric and monocentric systems. Genetics 232(1): iyaf247. doi: 10.1093/genetics/iyaf247.

Zhang H, Lang Z, Zhu J-K. 2018. Dynamics and function of DNA methylation in plants. Nat. Rev. Mol. Cell Biol.19: 489–506. doi: 10.1038/s41580-018-0016-z.

Zhang M, Steckenborn S, Castellani M, Marín-Gual L, Câmara AS, Robledillo LA, Maria-Parteka L, Sargheini N, Marek M, Chacón-Madrigal E, Gatica Arias A, Félix LP, Thomas WW, Huettel B, Pedrosa-Harand A, Lovell JT, Vanzela ALL, Ruiz-Herrera A, Marques A. 2026a. A holocentric pangenome links karyotype evolution to meiotic recombination. preprint, doi: 10.64898/2026.01.17.700048.

Zhang X, Hu Y, Huang K, Wright SI, Rieseberg LH. 2026b. Recombination suppression in plant adaptation and speciation. The New Phytologist 250: 2061–2077. doi: 10.1111/nph.71026.

