## Supplementary Figures S1-S12 for "Nowhere to Hide: The Effect of Centromere Architecture on LTR Retrotransposon Dynamics"

**Supplementary Figure S1. Assembly statistics for species included in the bioinformatic analysis.**

(**a**) Assembly size and haploid chromosome number across monocentric and holocentric species. Assembly size is shown on a log10-transformed scale, with axis labels back-transformed for readability. (**b**) BUSCO completeness assessed against the Viridiplantae odb10 conserved single-copy ortholog dataset and assembly gap density expressed as the number of gaps per Mbp. Marker color indicates plant family, and marker shape indicates centromere type. Most assemblies showed high completeness and low gap density, whereas *Oryza longistaminata* was identified as an outlier with elevated gap density and was excluded from further analyses.

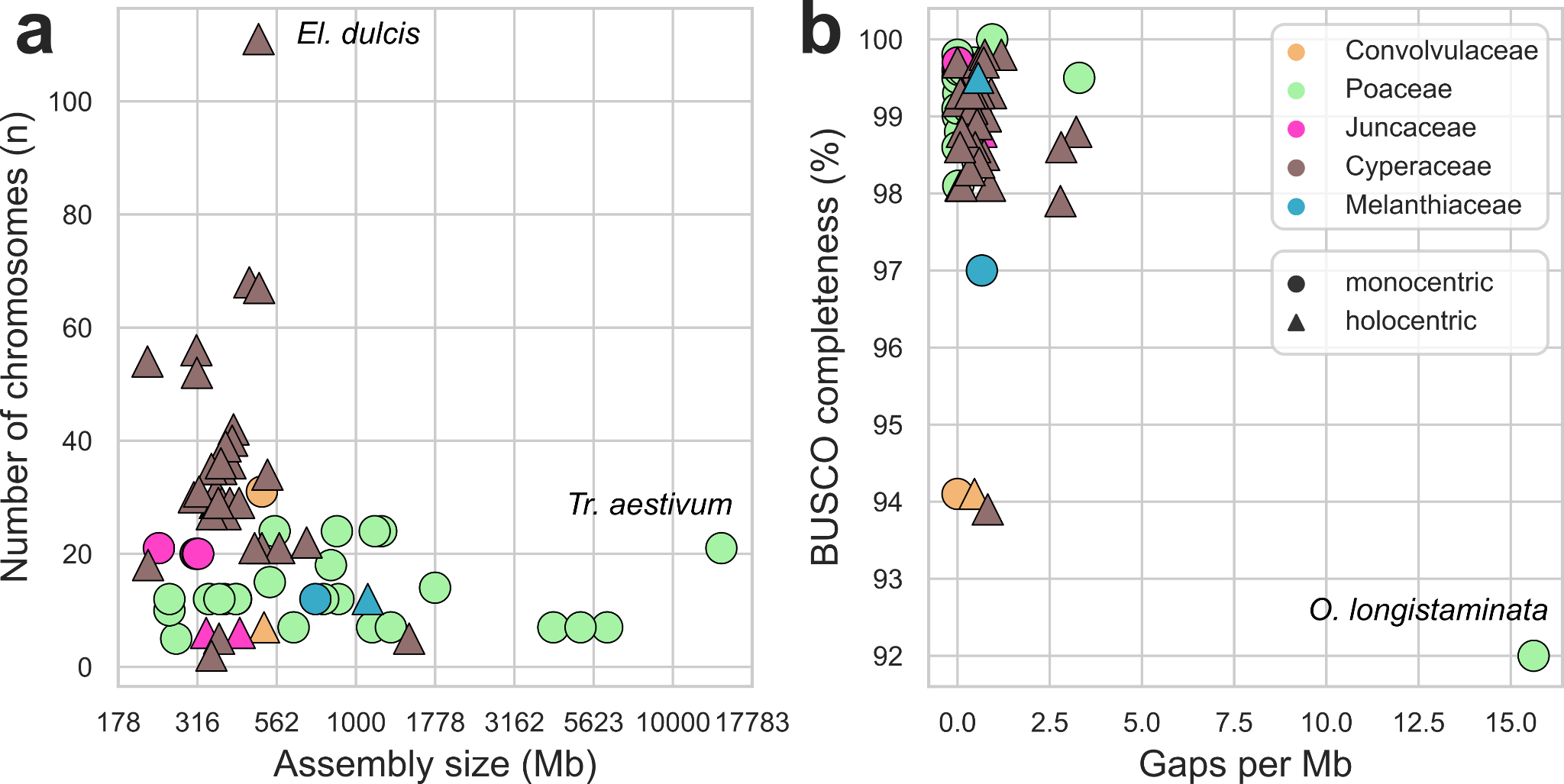

**Supplementary Figure S2. Chromosome-size effects on LTR-RT abundance and composition.**

Chromosome-level models of **(a,d)** LTR-RT density, **(b,e)** CRM proportion, and **(c,f)** Ty3-gypsy proportion in monocentric (grey) and holocentric (blue) chromosomes. Panels show analyses for the full dataset **(a-c)** and for the subset restricted to the chromosome-size range shared by both centromere types **(d-f)**.

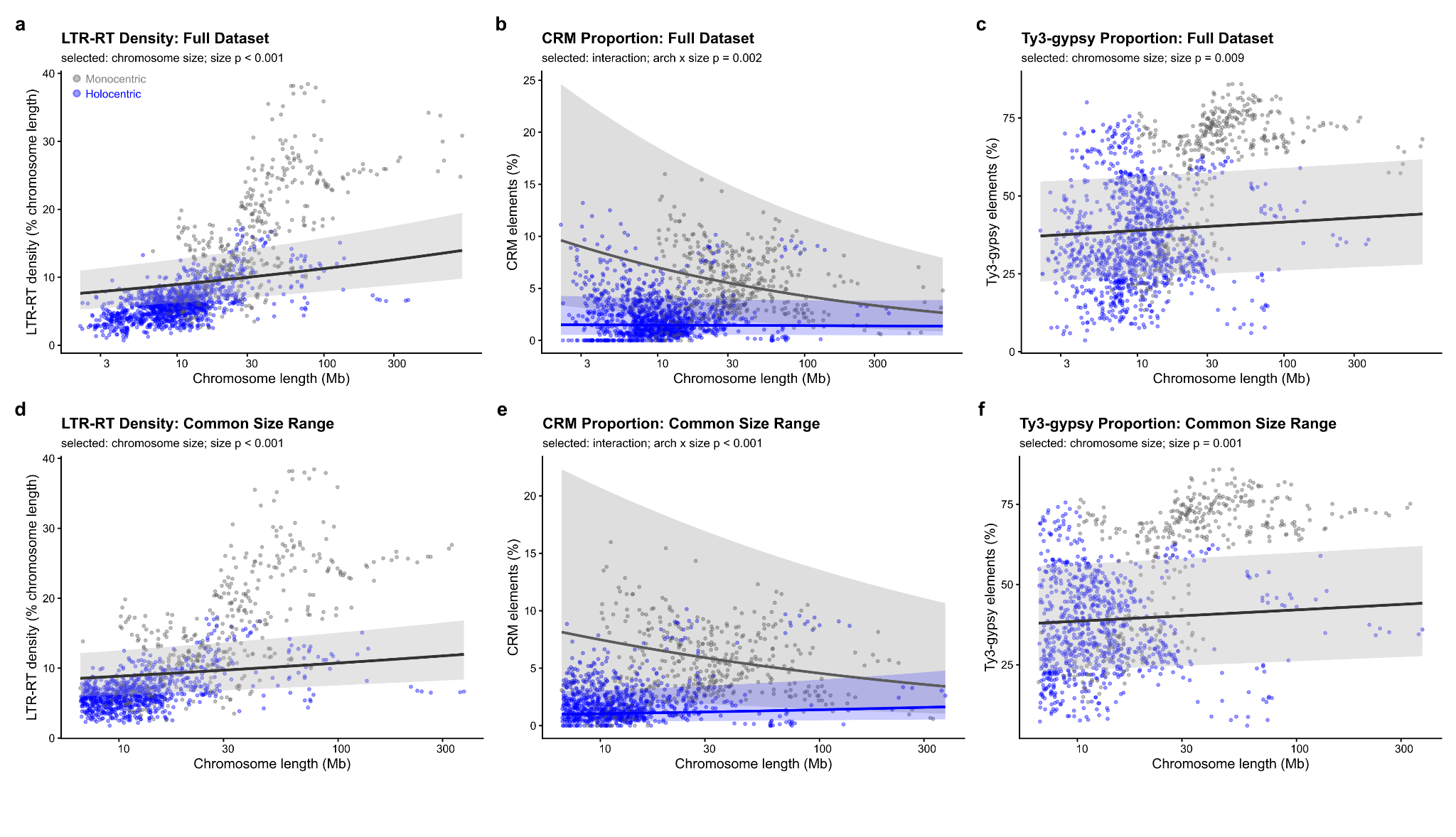

**Supplementary Figure S3. LTR identity in all LTR-RT families.**

Boxplots show the distribution of pairwise identity between 5′ and 3′ LTRs of full-length LTR retrotransposons for individual LTR-RT families across monocentric and holocentric species. LTR identity was used as a proxy for insertion age, with higher identity indicating more recent insertions. LTR identity values were logit-transformed prior to plotting to better distinguish differences near 100%. Axis tick labels are back-transformed for readability. Box boundaries represent the first and third quartiles, central lines indicate medians, and whiskers extend to values within 1.5× the interquartile range.

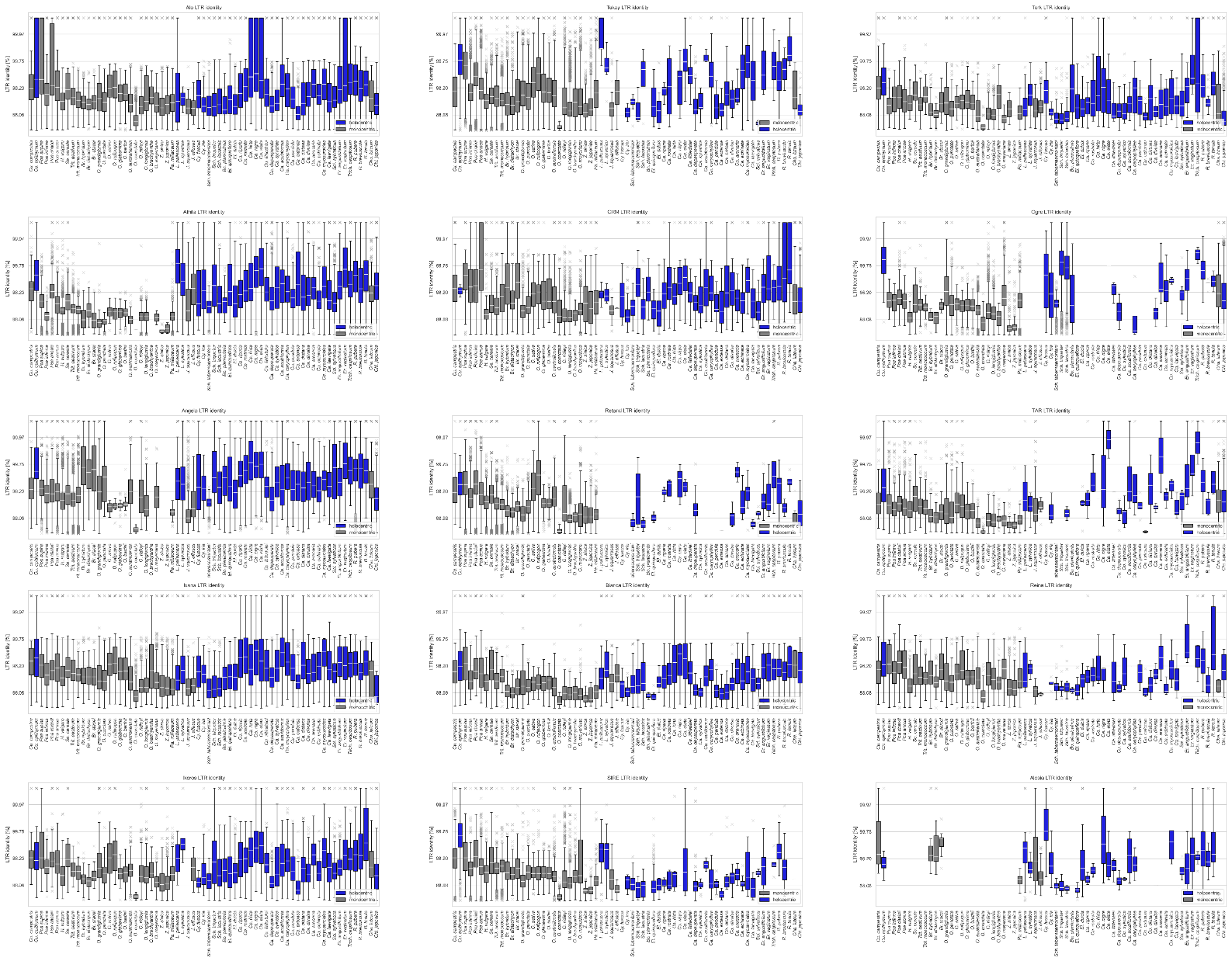

**Supplementary Figure S4. Scale-dependent spatial autocorrelation of age and LTR-RT removal signatures in monocentric and holocentric chromosomes.**

Distribution of Moran’s I values calculated from **(a)** genome-based LTR identity, **(b)** solo LTR to intact LTR retrotransposon ratios, and **(c)** FISH-based LTR to GAG/INT signal ratios across increasing neighborhood thresholds. Thresholds represent the proportion of chromosome length used to define neighboring windows. Asterisks indicate a significant effect of centromere architecture (monocentric vs. holocentric) on Moran's I at each threshold, from linear mixed models with architecture specified in **Supplementary Table S10** (*p < 0.05, **p < 0.01, ***p < 0.001). Unadjusted p-values are reported, since individual threshold values do not represent independent hypotheses.

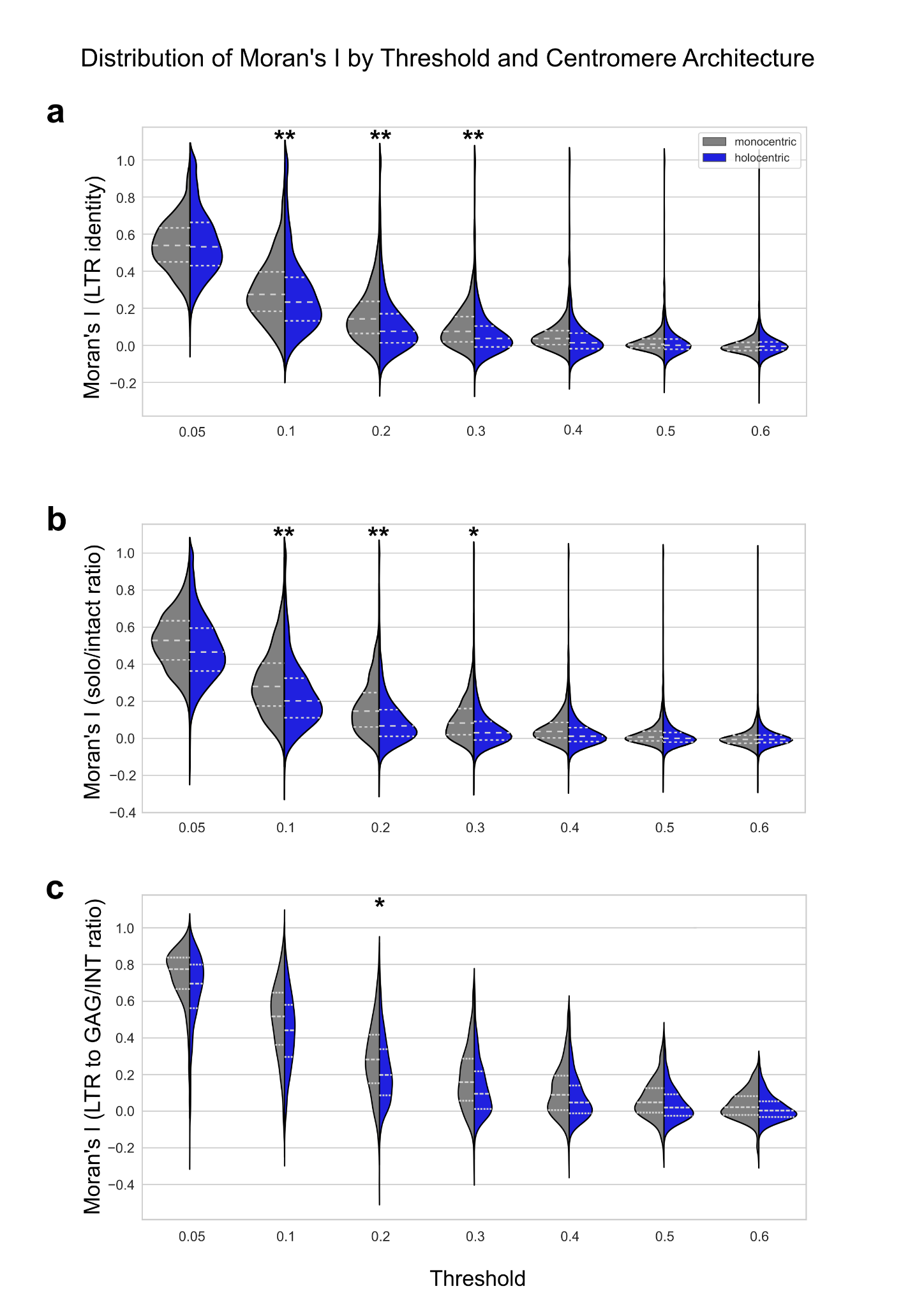

**Supplementary Figure S5. Dynamic Time Warping clustering of LTR identity and solo/full-length ratio profiles**

DTW clustering of **(a)** mean LTR identity profiles and **(b)** solo LTR to full-length LTR-RT ratio profiles along relative chromosome position. LTR identity profiles formed three archetypes (distal-elevated, uniform, and distal-depressed); solo/full-length ratio profiles formed two archetypes (distal removal and uniform removal). Bold lines show cluster mean profiles and thin lines show subsampled individual chromosome profiles. Dendrograms show hierarchical Ward clustering based on DTW distances. Stacked bar plots show archetype proportions in monocentric and holocentric species, split into quartiles according to chromosome size (1 - smallest, 4 - largest).

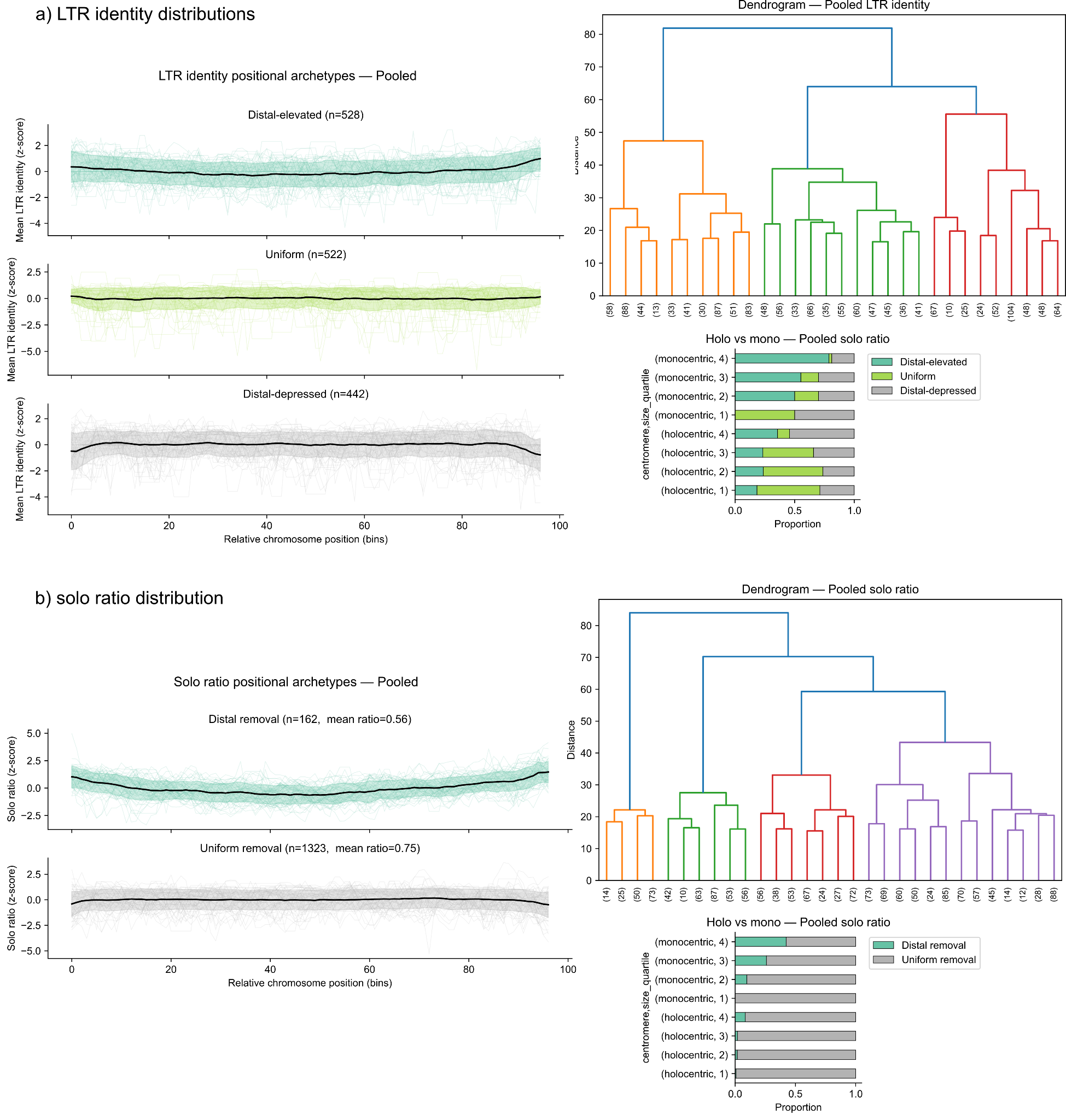

**Supplementary Figure S6. Comparison of genome-based and cytogenetic profiles of *Athila* LTR retrotransposons along *Hordeum vulgare* chromosome 6H.**

Chromosome 6H was identified in a metaphase spread using an rDNA-specific probe targeting the ITS1–ITS2 region. The genome-based profile shows the distribution of annotated *Athila* full-length elements, solo LTRs, and the ratio of solo LTRs to full-length elements along chromosome 6H. The cytogenetic profile shows FISH signal intensity along chromosome 6H using probes targeting *Athila* LTR and internal GAG regions, together with the LTR to GAG signal ratio. Both approaches reveal similar broad-scale patterns of *Athila* distribution along the chromosome.

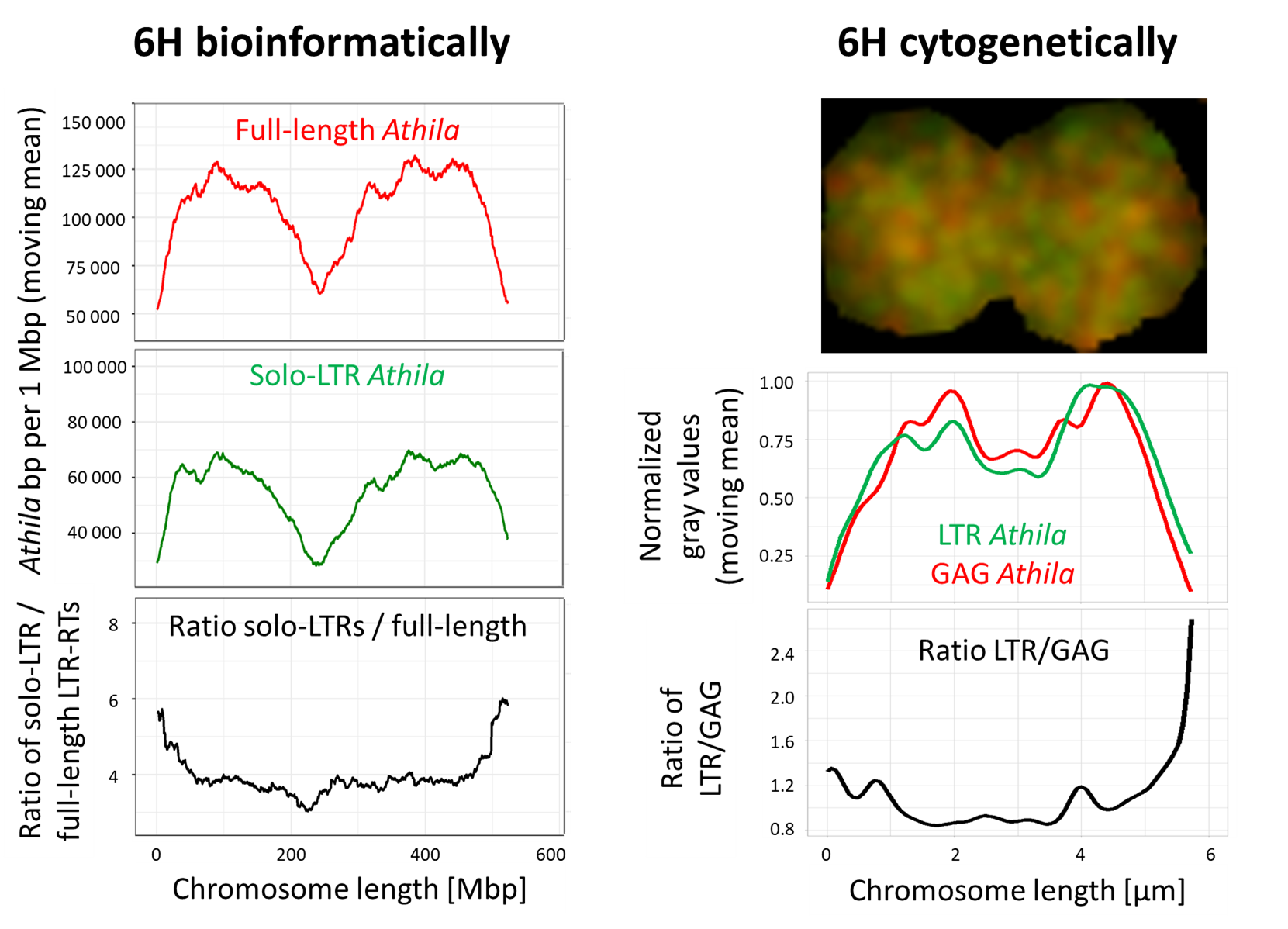

**Supplementary Figure S7. Family-specific FISH-based spatial autocorrelation of LTR retrotransposon signals.**

Moran’s I values calculated at a neighborhood threshold of 0.2 from FISH-based LTR to GAG/INT signal ratios for **(a)** *Angela* and **(b)** *Athila* family. The violin plots show the overall distribution of Moran’s I in monocentric and holocentric chromosomes, and the boxplots show Moran’s I distributions for individual species.

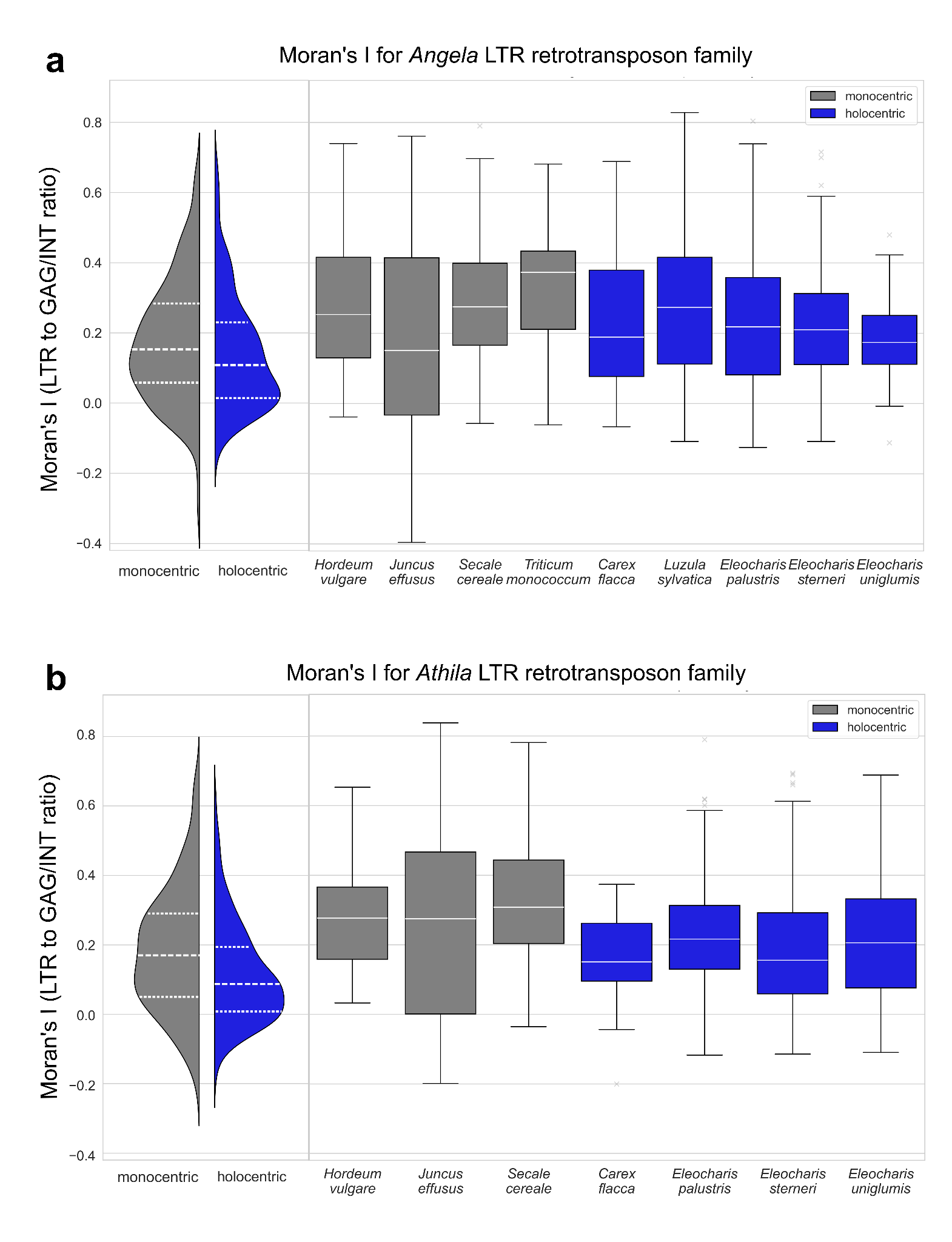

**Supplementary Figure S8. Sensitivity analysis of chromosome-size effects on LTR retrotransposon age, removal, and spatial organization in the shared chromosome-size range.**

This figure shows all LTR dynamics models in the chromosome-size range shared by monocentric and holocentric taxa. **(a)** LTR identity of intact LTR retrotransposons in relation to chromosome length restricted to the common chromosome-size subset. Higher LTR identity indicates more recent insertions. **(b)** Moran’s I of LTR identity in relation to chromosome length in the common chromosome-size subset. Higher values indicate stronger clustering of element age. **(c)** Window-level log-transformed solo/intact LTR-RT ratio in monocentric and holocentric chromosomes in the common chromosome-size subset. Partial residuals from the architecture-only sensitivity model are shown. **(d)** Moran’s I of the solo/intact LTR-RT ratio in monocentric and holocentric chromosomes in the common chromosome-size subset. Moran’s I values in **(b,d)** were calculated using a neighbourhood threshold of 0.2, corresponding to 20% of chromosome length. Grey denotes monocentric taxa and blue denotes holocentric taxa.

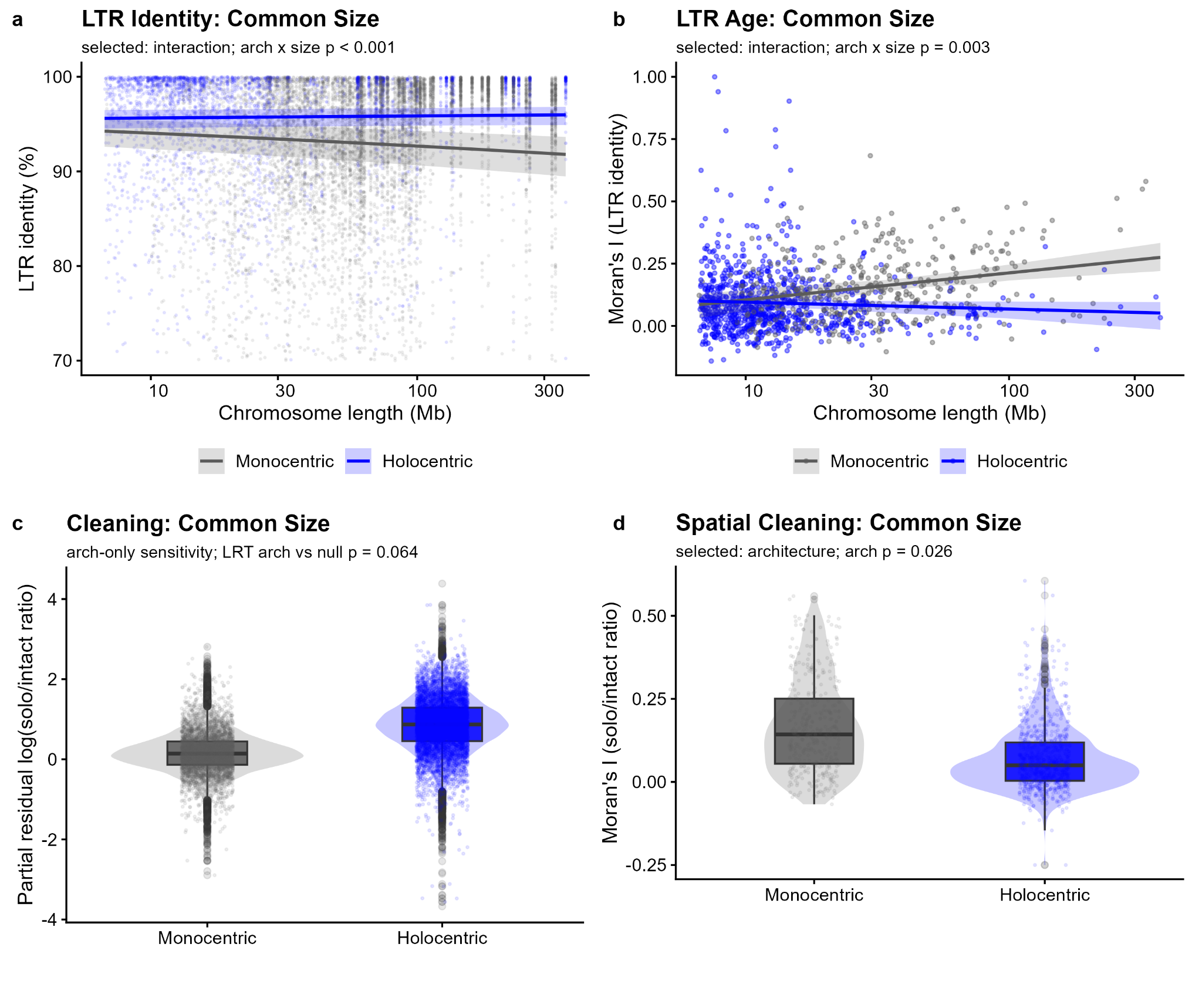

**Supplementary Figure S9. Correlation of genomic and epigenomic features in *Juncus effusus* and *Luzula sylvatica*.**

Spearman correlation matrices calculated in 100 kb genomic windows for monocentric *Juncus effusus* and holocentric *Luzula sylvatica*. Analyzed features include CenH3, histone modifications H3K9me2 and H3K4me3, DNA methylation in CpG, CHG, and CHH contexts, LTR retrotransposon categories (full-length intact elements, partial elements, and solo LTRs), LTR identity, gene density, and satellite repeat coverage. White cells indicate correlations that were not significant after Benjamini–Hochberg correction for multiple testing (FDR > 0.01).

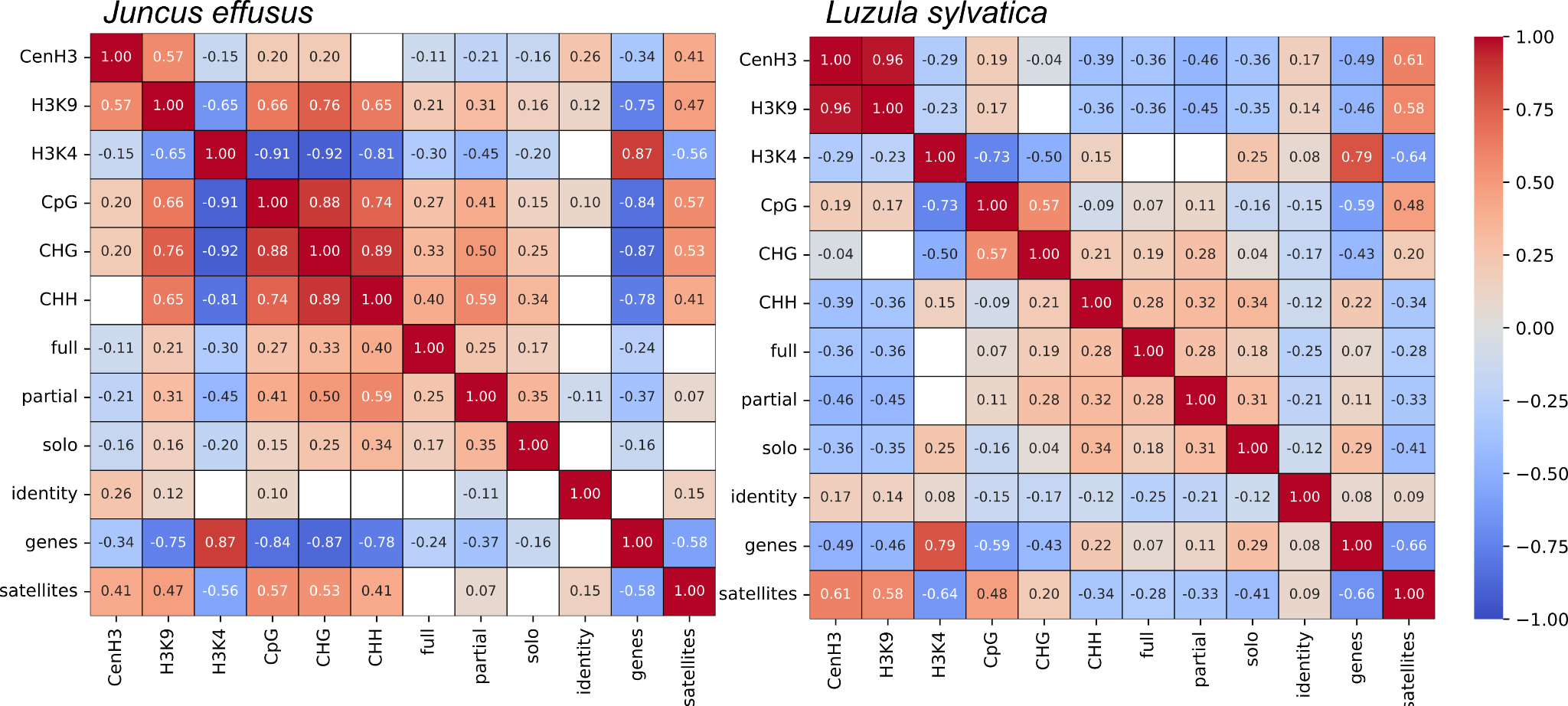

**Supplementary Figure S10. LTR identity of centromeric and non-centromeric LTR retrotransposons in *Juncus effusus* and *Luzula sylvatica*.**

Comparison of LTR identity between LTR retrotransposons overlapping CenH3-enriched centromeric regions and elements located outside these regions in monocentric *Juncus effusus* and holocentric *Luzula sylvatica*. LTR identity was used as a proxy for insertion age, with higher identity indicating more recent insertions. Centromeric elements show higher LTR identity than non-centromeric elements, consistent with the association of CenH3-enriched regions with young LTR retrotransposons.

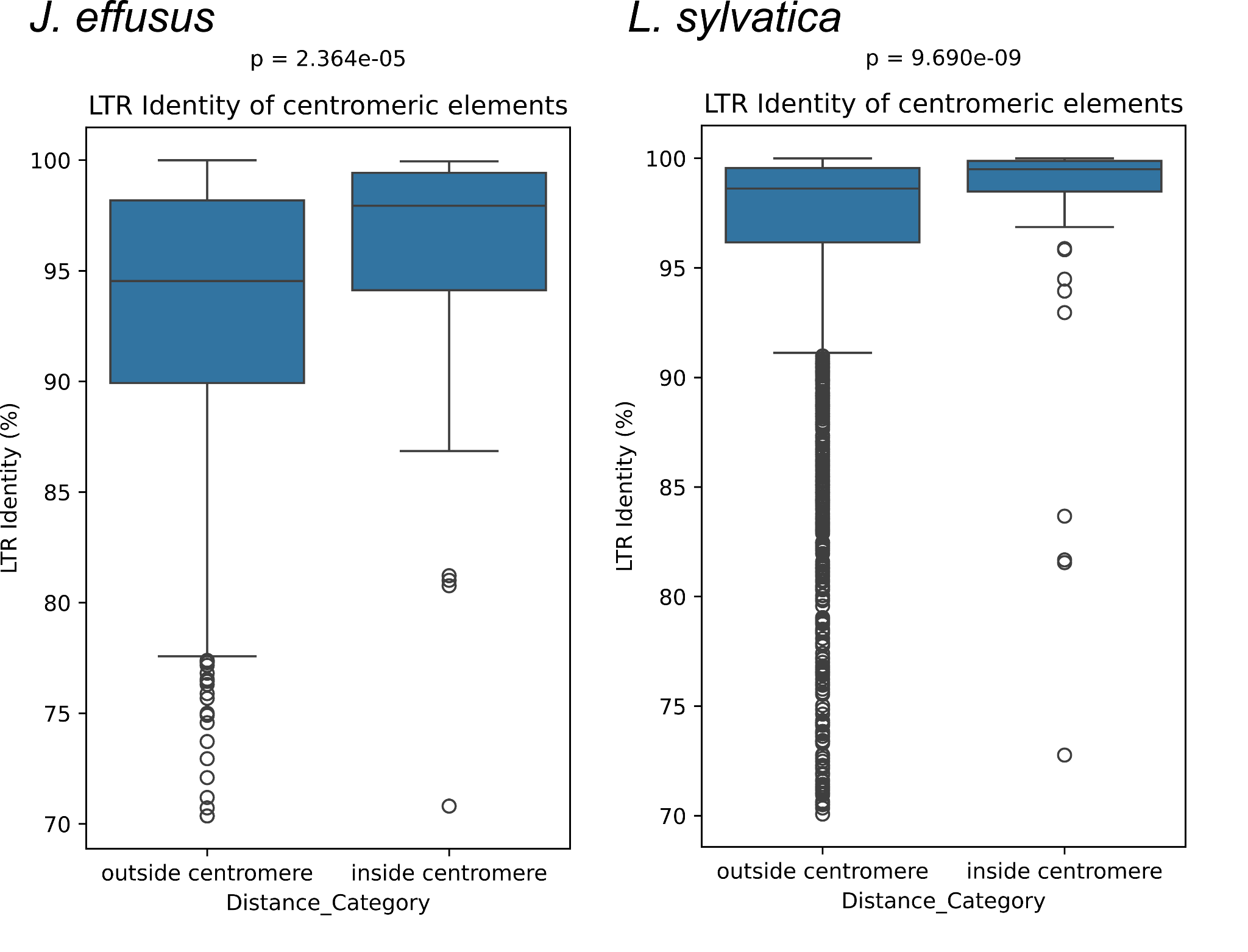

**Supplementary Figure S11. Epigenetic profiles of solo LTRs and LTRs within full-length LTR retrotransposons.**

Mean DNA methylation and ChIP-seq enrichment profiles of solo LTRs and LTR regions from full-length elements in *Juncus effusus* and *Luzula sylvatica*. Profiles are shown for random genomic regions (produced with *bedtools shuffle* command) and dominant LTR-RT families (*Ivana*, *Tork*, *Athila*, *Angela*) and include methylation in CpG, CHG, and CHH contexts, as well as CenH3, H3K9me2, and H3K4me3 enrichment. Solo LTRs show epigenetic profiles similar to LTRs within full-length elements, indicating that LTR-associated epigenetic regulation is largely retained after removal of the internal retrotransposon region.

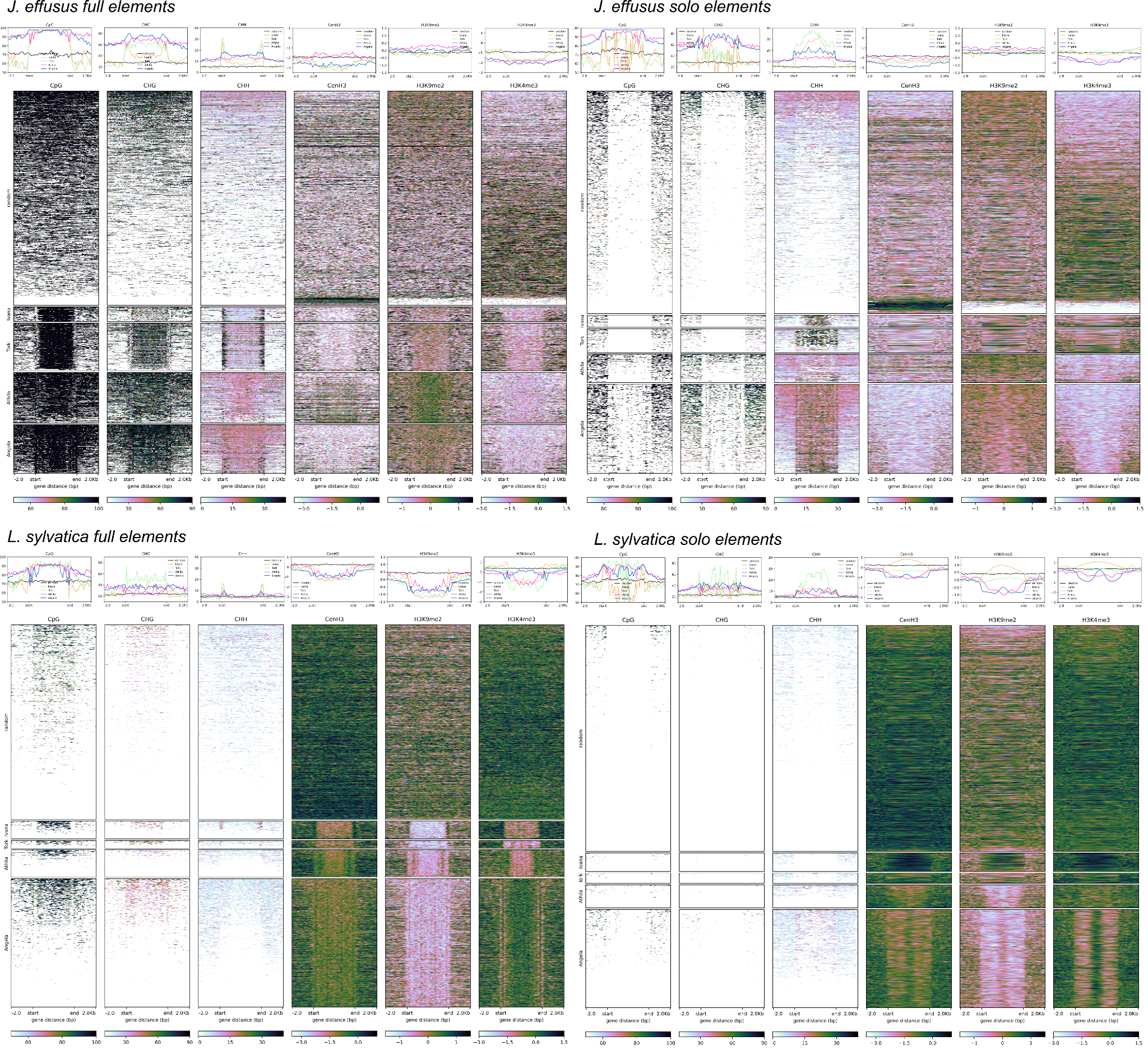

**Supplementary Figure S12. *Angela* LTR retrotransposon lineages in *Juncus effusus* and *Luzula sylvatica*.**

Pairwise distance heatmaps of reverse transcriptase sequences from *Angela* LTR retrotransposons in monocentric *Juncus effusus* and holocentric *Luzula sylvatica*. Reverse transcriptase sequences were clustered by sequence similarity, and the resulting distance matrices were visualized to compare lineage diversity and expansion patterns between the two species. The *Angela* family shows a pattern consistent with that of the *Athila* family shown in **Figure 5a**, with a dominant lineage in *L. sylvatica* and multiple co-dominant lineages in *J. effusus*.

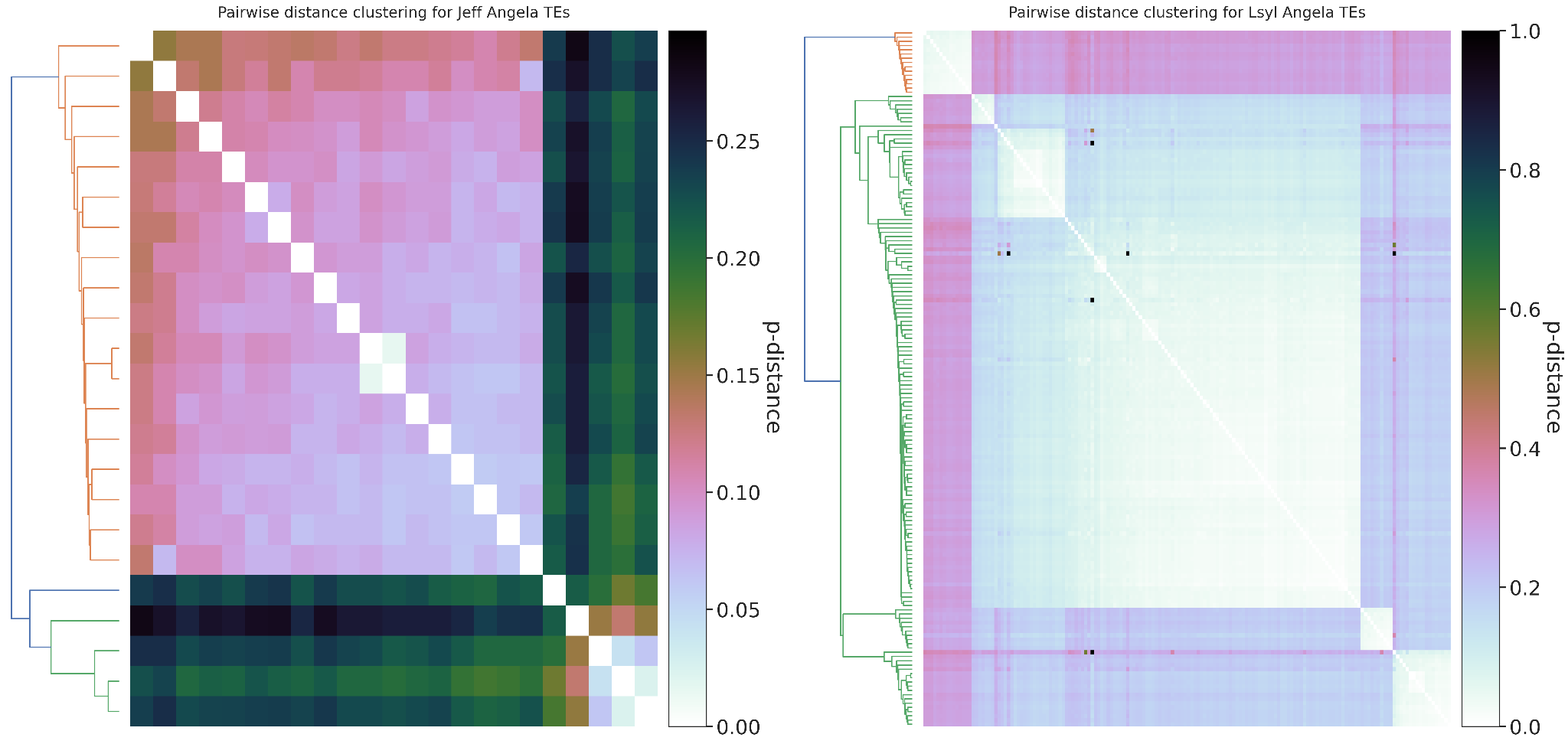
